# Precision CRISPR annotation of the functional enhancer landscape in primary human T cells

**DOI:** 10.64898/2026.01.27.702006

**Authors:** Ping Wang, Josephine R. Giles, Hua Huang, Sasikanth Manne, Martha S. Jordan, Alexander C. Huang, Junwei Shi, E. John Wherry

**Author notes:** Address correspondence to: J.R.G., J.S. and E.J.W.

## Abstract

Precise modulation of T cell function through engineering the non-coding genome holds great promise for advancing next-generation immunotherapies. However, robust high-throughput approaches to annotate functional cis-regulatory elements (CRE) in human T cells remain limited. Here, we developed a simple and highly efficient CRISPR interference (CRISPRi) perturbation platform to systematically annotate CREs in human primary T cells. Using this platform, we identified novel CREs controlling *PDCD1*, *HAVCR2*, and *TBX21* expression. Combinatorial CRE perturbations revealed synergistic CRE pairs that fine-tune *PDCD1* and *HAVCR2* expression, while Cas9-indel-based mutagenesis pinpointed the critical nucleotides within each enhancer that are essential for their activity. Functional experiments demonstrated that CRE-edited *HAVCR2* outperformed conventional total gene knockout in enhancing CAR T cell anti-tumor efficacy. Moreover, CRE editing of *PDCD1* and *HAVCR2* repressed PD-1 and TIM-3 expression in human tumor-infiltrating lymphocyte CD8 T cells, highlighting regulatory role of these CREs in disease relevant exhausted T cells. Together, this approach offers a compact CRISPRi platform that enables high-throughput dissection of functionally relevant non-coding genomic regions in T cells, providing insights for mechanistic studies and precision genome engineering of advanced cellular therapies.

## Main

T cell immunotherapy shows great promise for treating cancer and immune-related diseases^1,2^. Conventional genome engineering in primary human T cells has primarily focused on the protein-coding sequences, overlooking the 98%-99% of the genome where non-coding cis-regulatory elements (CREs), “enhancers” and “silencers”, reside. CREs play a critical role in controlling gene expression in a cell-type-specific manner, shaping cellular identity and function^3^. Manipulating CREs offers a nuanced strategy to precisely control gene expression in T cells for mechanistic studies and for cellular therapies. However, identifying functional CREs in primary human T cells remains challenging, restricting both our understanding of gene regulation and our capacity to optimize T cell function through engineering the regulatory landscape.

High-throughput epigenomic and transcriptomic profiling have identified hundreds of thousands to millions of candidate CREs genome-wide in human T cell subsets, but defining the functional relevance of these elements has remained challenging^4,5^. Developing unbiased functional genomic approaches to identify CREs in human T cells is essential to annotate the functional components of these transcriptional regulatory networks, and enable precise, cell-type-specific genome engineering. CRISPR-based approaches, including Cas9-mediated indel mutagenesis and CRISPR interference (CRISPRi), have enabled high-throughput screens of functional CREs at single-nucleotide or sub-kilobase resolution^6,7^. However, these screens have largely been confined to immortalized cell lines, due to low editing efficiency, challenges in transduction, and design constraints that restrict scalability in primary human T cells. This gap has hindered efforts to decode the gene regulatory networks that control human T cell function. Moreover, although CREs typically span hundreds of base pairs, only discrete short sequences within these DNA elements are essential for regulatory activity^8^. Mapping these functional elements in primary T cells is necessary to decode enhancer logic and enable precise genome engineering for immunotherapy.

Inhibitory receptors (IRs), such as PD-1, CTLA-4, LAG-3, and TIM-3, are major regulators of T cell responses. IRs maintain immune homeostasis by preventing excessive activation^9–12^, but in chronic disease they can suppress effective T cell activity and block the clearance of cancer cells or persistent pathogens, making these receptors (and their genes) attractive targets for immune engineering^13^. Indeed, targeting IRs, such as CTLA-4, PD-1, and LAG-3 has revolutionized treatments for human cancer in large part by modulating human T cell differentiation including reinvigorating exhausted T cells^14,15,16^. In preclinical models, other IRs including TIM-3 have received considerable attention for regulating T cell biology^17–19^. These strategies extend beyond cancer. For example, IR blockade or genetic ablation can enhance T cell anti-tumor activity, whereas agonistic therapies can suppress overactive immune responses in autoimmunity^20^. Thus, targeting IRs remains a central focus of therapeutic development across diverse diseases. Despite the success of targeting and ablating IR signaling clinically, these binary “on/off” interventions can disrupt immune balance, leading to adverse effects^21,22^. For example, although genetic deletion of PD-1 can initially enhance antiviral T cell responses during chronic infection, permanent loss of all PD-1 signals ultimately accelerates T cell exhaustion and causes these responses to decline over time^23^. In acute-resolving infection, PD-1 knockout (KO) CD8 T cells compromise the optimal formation of long-term memory^24^ and the differentiation of early “stem-like” CD8 T cell populations^9^. Indeed, in humans, full PD-1 KO in Chimeric Antigen Receptor (CAR) T cells fails to provide sustained benefit, and these cells are less fit over time^25,26^. These data highlight the need for more precise modulation of IR expression to achieve robust and durable therapeutic benefits. In mice, tuning PD-1 expression via deleting an enhancer specific to exhausted CD8 T cells improved CD8 T cell function and reduced immunopathology compared to full PD-1 genetic KO^27,28^. These observations highlight the potential of cell-state-specific enhancer targeting for tunable IR control in T cells. However, how human IRs and other key T cell differentiation genes are epigenetically regulated remains largely unexplored. Comprehensive functional interrogation of human T cell CREs is therefore essential to enable precise IR modulation, and such data may provide a foundation for advancing next-generation cell therapies via non-coding genome engineering.

Here, we developed a streamlined and optimized CRISPRi platform for systematic functional annotation of CREs in primary human CD8 T cells. By combining tiling and ATAC-seq-guided CRISPRi screens, we identified novel CREs regulating expression of key IRs, *PDCD1* (encoding PD-1) and *HAVCR2* (encoding TIM-3), as well as the transcription factor *TBX21* (encoding TBET) that controls effector genes. Saturation mutagenesis using Cas9-induced indels revealed critical DNA sequences within *PDCD1* and *HAVCR2* enhancers. Multiplexed CRE perturbations further uncovered CRE combinations that broadened the dynamic range of gene expression modulation. Functionally, editing of *HAVCR2* CREs outperformed conventional coding-region knockouts in a CAR T cell model. Moreover, CRE editing of *PDCD1* and *HAVCR2* effectively reduced PD-1 and TIM-3 expression in tumor-infiltrating lymphocyte (TIL) CD8 T cells. Altogether, our study presents a compact CRISPRi platform that enables high-throughput functional dissection of non-coding regions in primary human T cells, offering insights for gene regulation and opportunities for precision genome engineering and improved cellular therapies.

## Results

### Development of a high-efficiency all-in-one CRISPRi platform for CRE screening in primary human CD8 T cells

We set out to optimize a CRISPRi system to enable high-throughput functional annotation of CREs in primary human T cells. Traditional CRISPRi relies on a catalytically dead Cas9 (dCas9) protein fused to a wild-type Krüppel-associated box (KRAB) domain, to recruit repressive complexes and deposit H3K9me3 on chromatin, enabling CRE scanning at ∼0.5-2 kb resolution^29–31^. Previous work employed a dual-vector lentiviral system, one encoding the sgRNA and another encoding the dCas9-KRAB, to target CREs in primary T cells^32^. This approach suffered from low co-transduction efficiency, limiting scalability to large-scale screens. To address this limitation, we engineered an all-in-one CRISPRi system by combining the sgRNA and dCas9-KRAB cassettes (driven by bidirectional promoters) into a single lentiviral vector co-expressing a GFP marker, enabling tracking of CRISPRi positive cells (Extended Data Fig. 1a). This system achieved an average CRISPRi transduction efficiency of 33.3% in primary human CD8 T cells, compared to 12.2% with the conventional dual-vector system (Extended Data Fig. 1b, c). To further enhance silencing efficiency, we tested two engineered variants of the ZNF10-KRAB domain (WSR8EEE-KRAB and AW7EE-KRAB)^33^ and one KRAB from ZIM3^34^ using sgRNAs targeting of the *CXCR3* promoter. CXCR3 is a chemokine receptor expressed widely by memory CD8 T cells in human blood and can be easily measured by flow cytometry. Surface expression of CXCR3 was measured by flow cytometry as a proxy for gene silencing. All three alternative KRAB variants outperformed wild-type ZNF10-KRAB, with ZIM3-KRAB exhibiting the highest silencing efficiency, increasing repression from ∼69% to ∼85% (Extended Data Fig. 1d, e).

Next, we designed a pooled sgRNA library screen to evaluate the performance of the improved all-in-one CRISPRi system (Fig. 1a). We selected the *CXCR3* genomic region as a model to assess the capacity of this system for identifying functional CREs, given the relative simplicity of the *CXCR3* locus and the ease of monitoring cell surface protein expression^35^. Two sgRNA libraries were designed: (1) an accessible chromatin region (ACR)-guided library consisting of 231 sgRNAs targeting 17 ACRs identified by ATAC-seq in primary human T cells, and (2) a tiled library of 901 sgRNAs spaced at 200bp intervals spanning ∼292kb upstream and ∼42kb downstream of the *CXCR3* locus (for a total of a ∼335kb genomic region; Supplementary Table 2). Both libraries included positive control sgRNAs targeting the *CXCR3* promoter and a previously validated enhancer^32^, along with non-targeting sgRNAs and sgRNAs for non-accessible unrelated genomic regions as negative controls. Human primary CD8 T cells from two healthy donors were activated *in vitro* and transduced with each pooled library at low multiplicity of infection. On day 5 post transduction, cells in the top and bottom 20% of CXCR3 surface expression were isolated at a minimum of 1000 cells/sgRNA coverage of the library, and sgRNA representation was quantified by deep sequencing. To assess the performance of each sgRNA, a CRISPRi score was calculated as the log2-fold change (Log2FC) in abundance of the bottom versus the top bins (Fig. 1a). For the tiled library, we leveraged the high sgRNA density by applying a sliding-window analysis. Using an optimized 20-sgRNA window, we averaged consecutive sgRNA scores to minimize variability of individual sgRNAs, enhancing peak detection and capturing regional regulatory context with the highest inter-donor correlation (Extended Data Fig. 2a, b). Results from two donors were well correlated in both the ACR and tiling screens (Extended Data Fig. 3a), and the negative control sgRNAs centered around 0, indicating low noise in the pooled library screens (Extended Data Fig. 3b).

**Fig. 1:**
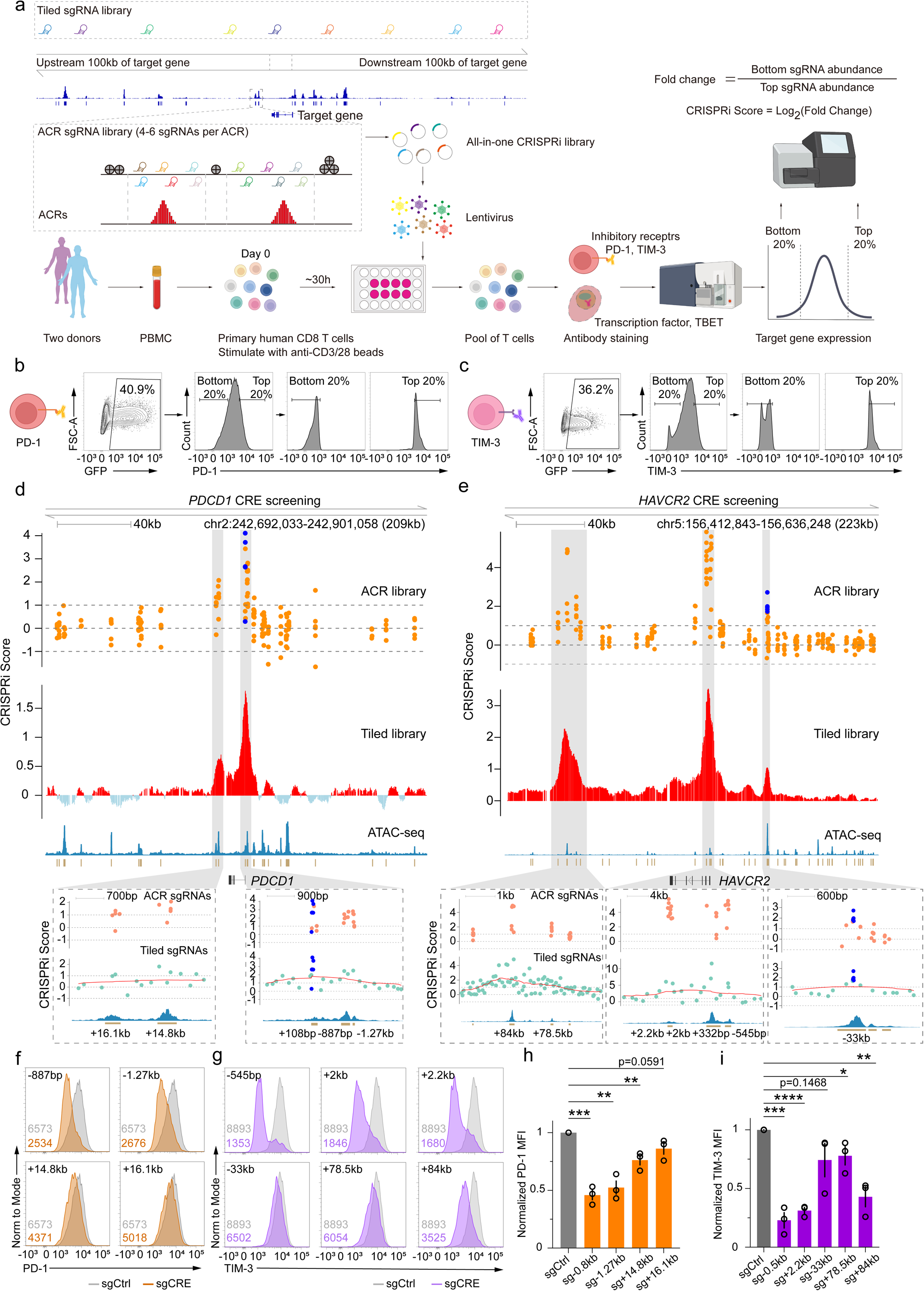
All-in-one CRISPRi screening platform identified functional CREs of *PDCD1* and *HAVCR2*. a, Schematic of all-in-one CRISPRi pooled screens for discovering functional CREs in primary human CD8 T cells. Two parallel sgRNA libraries were designed: library 1 targeted ACRs within 100kb upstream and downstream of the TSS with 4-6 sgRNAs designed per ACR; library 2 tiled this region with sgRNAs at 200bp intervals. **b,c,** Representative flow cytometry plots showing sorting of the top and bottom 20% of cells based on PD-1 (**b**) or TIM-3 (**c**) expression for the pooled CRISPRi genetic screening. **d,e,** Results of pooled CRISPRi screens for CRE of *PDCD1* (**d**) or *HAVCR2* (**e**) using ACR and tiled library designs (donor ND584). Top, ACR library screen. Blue dots, pre-validated positive controls. Orange dots, sgRNAs targeting ACRs. Middle, tiled library screen. Red and light blue bars indicate average CRISPRi scores for the tiled library screening, calculated based on a sliding window of 20 consecutive sgRNAs. Bottom, ATAC-seq profiles of the corresponding genomic regions. Gray shaded areas are zoomed-in regions with high CRISPRi scoring sgRNAs (Log2FoldChange > 1). Green dots, sgRNAs from the tiled library. Red curves represent the same data from the red bars as a smoothed line. **f,g**, Representative flow cytometry plots showing validation of putative CREs using individual sgRNAs, CRISPRi-mediated reduction of PD-1(**f**) or TIM-3 (**g**) expression. Numbers represent MFI, colors coded accordingly. **h,i**, Quantification of MFI from (**f**) and (**g**) for CRISPRi-mediated reduction of PD-1 (**h**) and TIM-3 (**i**). Data represent mean ± s.e.m.; n = 3 donors, three independent experiments. *P < 0.05, **P < 0.01, ***P < 0.001, ****P<0.0001, two-tailed Student’s t-test.

The screening libraries identified the previously defined enhancer as well as two additional proximal CREs. Across all screens, sgRNAs targeting regions proximal to the TSS yielded the strongest silencing of *CXCR3* (Extended Data Fig. 3c-f). The previously described -4.1kb enhancer of *CXCR3* ^32^ was also re-identified using independent sgRNAs from both libraries, confirming the robustness of this screening approach (Extended Data Fig. 3c-f). Two new CREs located 1.6 kb and 2.3 kb downstream of the *CXCR3* TSS were identified, showing strong CRISPRi scores above background (Extended Data Fig. 3c-f). This +2.3 kb CRE was not included in the ACR library because this region had variable chromatin accessibility across multiple donors in a reference ATAC-seq dataset^32^. This region scored highly, however, in the tiled library in both donors tested in this screen. To validate the screening results, the candidate +1.6 kb, +2.3 kb, and -4.1 kb CREs were individually tested. Targeting each of these three regions resulted in strong reduction of CXCR3 protein expression measured by flow cytometry (Extended Data Fig. 3g). Together, these results demonstrate the high efficiency and low noise of our optimized all-in-one CRISPRi system to discover known and novel functional regulatory elements in human primary CD8 T cells.

### All-in-one CRISPRi screening identifies functional CREs of inhibitory receptors in human primary CD8 T cells

Engineering the non-coding genome offers a powerful and precise way to modulate gene expression. Tuning expression of IRs by T cells could be used to restore T cell function during chronic infection and cancer, while limiting risks of immune pathology. To this end, we performed unbiased genetic screens targeting the genomic regions surrounding *PDCD1* (encoding PD-1) and *HAVCR2* (encoding TIM-3), two key IRs involved in regulating CD8 T cell responses in cancer and chronic infections. First, we performed a time-course analysis of PD-1 and TIM-3 expression in CD8 T cells from two healthy donors to determine the optimal time points for CRISPRi-mediated repression. Flow cytometry measurement after sgRNAs targeting the IR promoters revealed days 4 and 6 post-activation as the optimal timepoints for detecting surface PD-1 and TIM-3 repression, respectively (Extended Data Fig. 4a, b). We designed both ACR and tiling sgRNA libraries targeting ∼200 kb flanking regions (+100kb and -100kb) of *PDCD1* or *HAVCR2* genomic loci. The tiling libraries contained over 1,000 sgRNAs and the ACR libraries included 200-300 sgRNAs (Supplementary Tables 3 and 4). Screens were performed in CD8 T cells from two donors as described in the *CXCR3* screens (Fig. 1a). The top and bottom 20% of cells expressing the protein of interest were sorted at the peak repression timepoints (day 4 for *PDCD1* and day 6 for *HAVCR2*) and subjected to deep sequencing to determine the CRISPRi scores (Fig. 1a-c). Results for *PDCD1* and *HAVCR2* were well correlated between donors for both libraries (Extended Data Fig. 5a, b).

ACR and tiling screens identified candidate functional CREs for *PDCD1* and *HAVCR2*. Negative controls and promoter-targeting sgRNAs behaved as expected (Fig. 1d, e; Extended Data Fig. 5c, d). The tiling screen uncovered distinct distal CRE peaks, including previously uncharacterized elements approximately 15 kb downstream of the *PDCD1* TSS, as well as −33 kb and +80 kb from the *HAVCR2* TSS (Fig. 1d, e; Extended Data Fig. 5c, d). Guided by the ACR screens, we found focal enrichment of multiple active sgRNAs at putative CREs: +14.8 kb and +16.1 kb for *PDCD1*, and −33 kb, +78.5 kb, and +84 kb for *HAVCR2* (Fig. 1d, e; Extended Data Fig. 5c, d).

To validate the screening results, the newly discovered CREs were individually tested by CRISPRi using three additional donors. Flow cytometry revealed a 15-25% reduction in PD-1 surface expression for sgRNAs targeting the *PDCD1* +14.8kb and +16.1kb CREs, compared to ∼50% reduction with a promoter-targeting sgRNA (Fig. 1f, h). Similarly, sgRNAs targeting *HAVCR2* CREs located at -33 kb, +78.5 kb, and +84 kb decreased TIM-3 expression by 25-50%, versus a 75% reduction with TSS-targeting sgRNAs (Fig. 1g, i). qPCR analysis confirmed corresponding reductions in *PDCD1* and *HAVCR2* mRNA expression, consistent with the protein expression changes and the original screening results (Extended Data Fig. 5e, f).

To further evaluate the enhancer activity of these distal CREs, we employed a minimal promoter-driven GFP reporter system. Full-length DNA fragments of *PDCD1* +16.1kb CRE and the three distal *HAVCR2* CREs (−33 kb, +78.5 kb, and +84 kb) were individually cloned into lentiviral reporter constructs and transduced into primary human CD8 T cells (Extended Data Fig. 6a). The *PDCD1* +16.1 kb CRE increased GFP expression ∼2.6-fold, whereas the *HAVCR2* CREs (−33 kb, +78.5 kb, +84 kb) increased GFP by 2.5–6-fold compared to negative controls (Extended Data Fig. 6b,c). Altogether, these data indicate that this all-in-one CRISPRi platform is capable of identifying novel functional CREs that regulate *PDCD1* and *HAVCR2* expression in primary human CD8 T cells.

### All-in-one CRISPRi screening identifies functional CREs of *TBX21* in primary human CD8 T cells

IRs are of considerable interest for controlling T cell function and exhaustion and can be easily screened in the approaches above because of cell surface protein expression. Transcription factors (TF) also have a major role in regulating T cell function and differentiation, often governing expression of suites of related genes and gene programs. It was unclear, however, whether an intracellular staining approach to assess changes in protein expression would be compatible with the sequencing readouts needed for the CRE screening. To address this question and test the broader applicability of genetic screening beyond cell-surface proteins (e.g. IRs), we evaluated the performance of our all-in-one CRISPRi system for studying CREs regulating TFs in T cells, where measuring protein expression changes requires cell fixation and permeabilization. We focused on *TBX21* (encoding TBET), a master TF that enhances IFN-γ production and the effector T cell cytotoxicity program^36^. Because of fluorescent signal loss caused by cell fixation and permeabilization during intracellular staining, we replaced the GFP reporter with Thy1.1 (CD90.1), a congenic cell-surface protein that can be stained and used to sort CRISPRi positive cells. Given the correlation between ACR and tiling library screens in the studies above, we designed only an ACR library for *TBX21*, consisting of 350 sgRNAs, targeting 40 ACRs across a 213kb genomic region flanking *TBX21* (Supplementary Table 5). Pooled CRISPRi screens were performed in CD8 T cells from two donors, with cells sorted into the top and bottom 20% of TBET-expressing populations 5 days post-activation, followed by sequencing (Fig. 2a and Extended Data Fig. 4b).

**Fig. 2:**
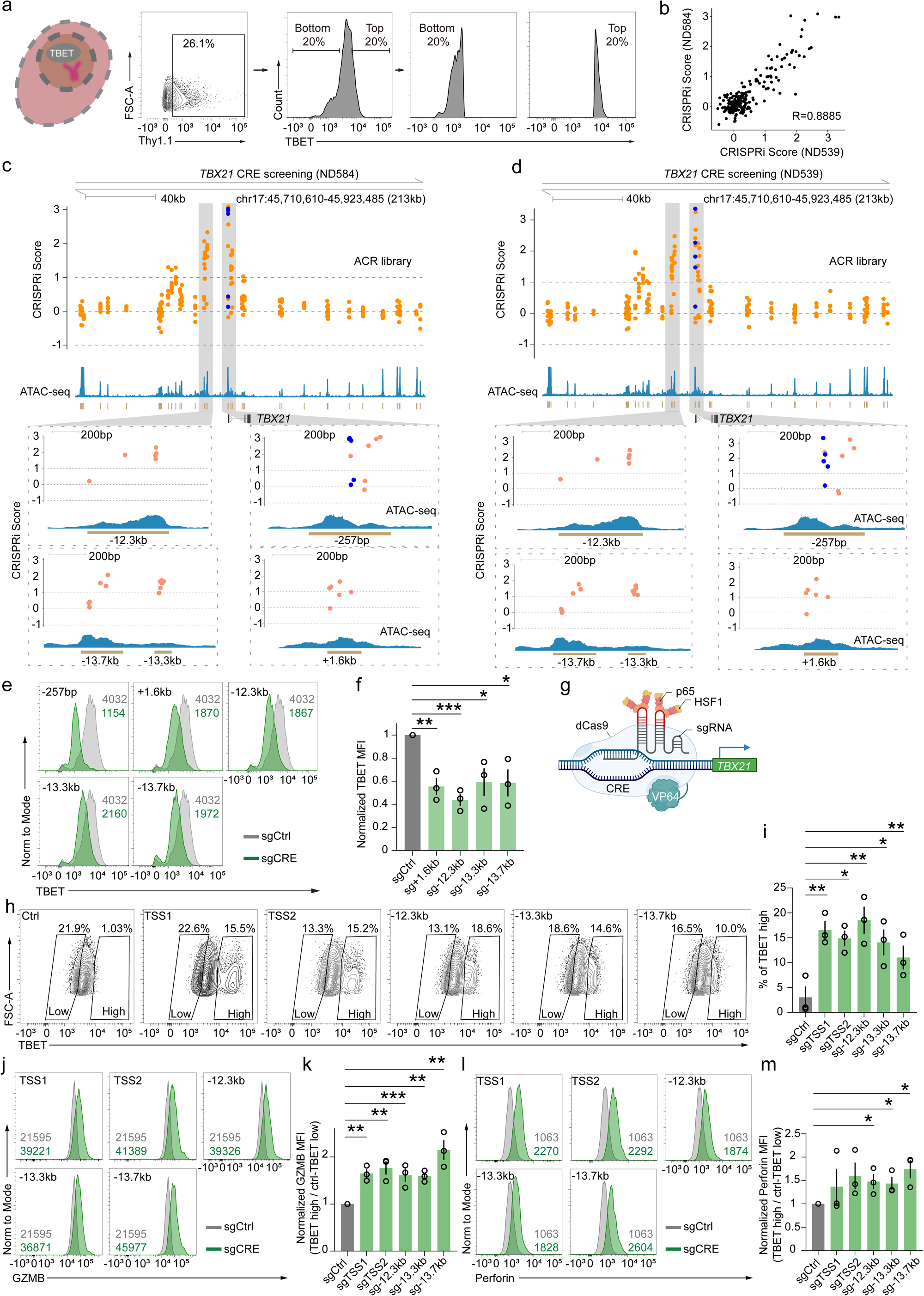
All-in-one CRISPRi screening platform identified functional CREs regulating *TBX21* expression. a, Flow cytometry plots showing sorting of the top and bottom 20% of cells based on TBET expression for the pooled CRISPRi genetic screening. **b,** Scatter plot showing reproducibility of CRISPRi screening across two donors. Each dot represents a single sgRNA (R = 0.8885, Pearson correlation). **c,d**, ACR library screens for functional CREs of *TBX21* in donor ND584 (**c**) and donor ND539 (**d**). Blue dots denote positive controls. Orange dots represent sgRNAs targeting ACRs. Gray shaded areas are zoomed-in regions containing high CRISPRi score sgRNAs (Log2FoldChange>1). **e**, Representative flow cytometry plots of TBET expression in negative control, TSS, and CRE targeted T cells by CRISPRi (numbers on top right indicate MFI). **f**, Quantification of TBET expression from (**e**) across donors. Data represent mean ± s.e.m.; n=3 donors, three independent experiments. **g,** Schematic of using CRISPRa to activate *TBX21*. **h,** Representative flow cytometry plots of TBET expression in negative control, *TBX21* TSS and CRE targeted groups by CRISPRa. **i**, Summary of TBET high proportion quantified from (**h**) (n=3 donors, mean ± s.e.m.; two independent experiments). **j,** Representative flow cytometry histograms of GZMB expression in TBET high cells with *TBX21* TSS or CREs targeted by CRISPRa (numbers on bottom left indicate MFI). **k,** Quantification of GZMB expression from (**j**) across donors (n=3 donors, mean ± s.e.m.; two independent experiments). **l,** Representative flow cytometry histograms of Perforin expression in TBET high cells with *TBX21* TSS or CREs targeted by CRISPRa (numbers on bottom right indicate MFI). **m,** Quantification of Perforin expression from (**l**) across donors (n = 3 donors, mean ± s.e.m.; two independent experiments). *P < 0.05, **P < 0.01, ***P < 0.001, two-tailed Student’s t-test. Numbers for MFIs are color coded accordingly.

This TF-focused CRE screening approach produced robust signals and enabled the identification of CREs controlling TBET expression and the results between donors were well correlated (Fig. 2b). Positive control sgRNAs targeting the TSS (−257bp and +1.6kb) were readily recovered in the screen (Fig. 2c, d). Three additional distal regions, located at -12.3kb, -13.3kb, and -13.7kb, were also identified as candidate CREs of *TBX21* (Fig. 2c, d). To verify the function of putative CREs, we selected high-scoring sgRNAs to target each CRE individually. qRT-PCR and flow cytometry confirmed that targeting these regions reduced *TBX21* mRNA and TBET protein expression to varying degrees, consistent with screening results, respectively (Extended Data Fig. 5g; Fig. 2e, f). Using the minimal promoter-driven GFP reporter system described above, we further demonstrated that inclusion of full-length DNA fragments of these three CREs increased GFP expression over 2-fold compared to the control (Extended Data Fig. 6b, c). TBET is a master regulator of effector gene expression in T cells including GZMB and Perforin^37^. We, therefore, next employed CRISPR activation (CRISPRa) to investigate the role of the identified CREs in regulating these cytotoxic genes (Fig. 2g). Targeting the *TBX21* TSS or CREs generated TBET high cells whereas there was little TBET expression in the negative control sgRNA cells (Fig. 2h). TSS targeting resulted in 15%-20% TBET high populations compared to the negative control, and targeting the CRE yielded 10%-20% TBET high cells (Fig. 2i). Thus, this CRE CRISPR screening platform was capable of being adapted for assessment of regulatory regions even for proteins that required intracellular staining.

One motivation for adapting this screening strategy to intracellular proteins was to probe the regulatory control of TFs. Given that TFs often sit at the apex of transcriptional circuitry controlling coordinated gene programs, we reasoned that targeting CREs in the *TBX21* locus regulating TBET expression would elicit corresponding changes in other TBET-dependent genes. Indeed, GZMB and Perforin expression were significantly higher in cells where CRISPRa targeting of *TBX21* CREs led to higher TBET expression, compared to TBET-low cells from the negative control. In these cells, there was a 1.6–2.2 fold increase in GZMB and a 1.4–1.7 fold increase in Perforin expression (Fig. 2j–m), demonstrating that CRE-mediated enhancement of TBET expression was both transcriptionally and biologically functional. Collectively, these data identify novel CREs and highlight the utility of this all-in-one CRISPRi platform for high-throughput CRE screening of genes encoding not only cell-surface proteins, but also intracellular molecules such as TFs.

### Fine-mapping of critical sequences in CREs using Cas9-mediated indel saturation mutagenesis

CREs control gene expression through multiple mechanisms including the binding of regulatory factors, such as TFs, at specific DNA sequences. Whereas CRISPRi-based perturbations can identify functional CREs at sub-kilobase resolution, dissecting these specific short DNA sequences underlying CRE activity requires higher-resolution perturbations. CRISPR-Cas9-induced double-strand breaks (DSB) at critical sites within CREs can disrupt their activity, enabling high-resolution dissection of CREs^8^. To pinpoint the specific sequences that confer function of the *PDCD1* and *HAVCR2* CREs, we used a Cas9-induced saturation mutagenesis strategy. This approach creates small insertions or deletions (typically 1–10 bp) at target sites within these CREs and enables screening with densely tiled sgRNAs libraries spanning each enhancer.

To enable Cas9-indel screening in a single-vector format, we modified our all-in-one CRISPRi construct by replacing dCas9-ZIM3 with catalytically active Cas9. All possible sgRNAs were designed to target the two novel CREs of *PDCD1* and the three CREs of *HAVCR2* identified from our CRISPRi screens (∼190 sgRNA for each; Supplementary Table 6). For each library, positive controls included guides targeting exons and the 1kb upstream region of each TSS. Negative control sgRNAs were from the previous CRISPRi library and newly designed guides targeting 100 bp flanking sequences on either side of each enhancer. Screens were performed in CD8 T cells from two donors and CRISPR scores were generated for the top and bottom 20% of *PDCD1* or *HAVCR2* expressing cells, following the same workflow as our CRISPRi screens (Fig. 3a). Screens targeting *HAVCR2* CREs were highly consistent across donors, while at the *PDCD1* CREs showed only modest concordance, indicating potential donor-specific effects (Extended Data Fig. 7a, b).

**Fig. 3:**
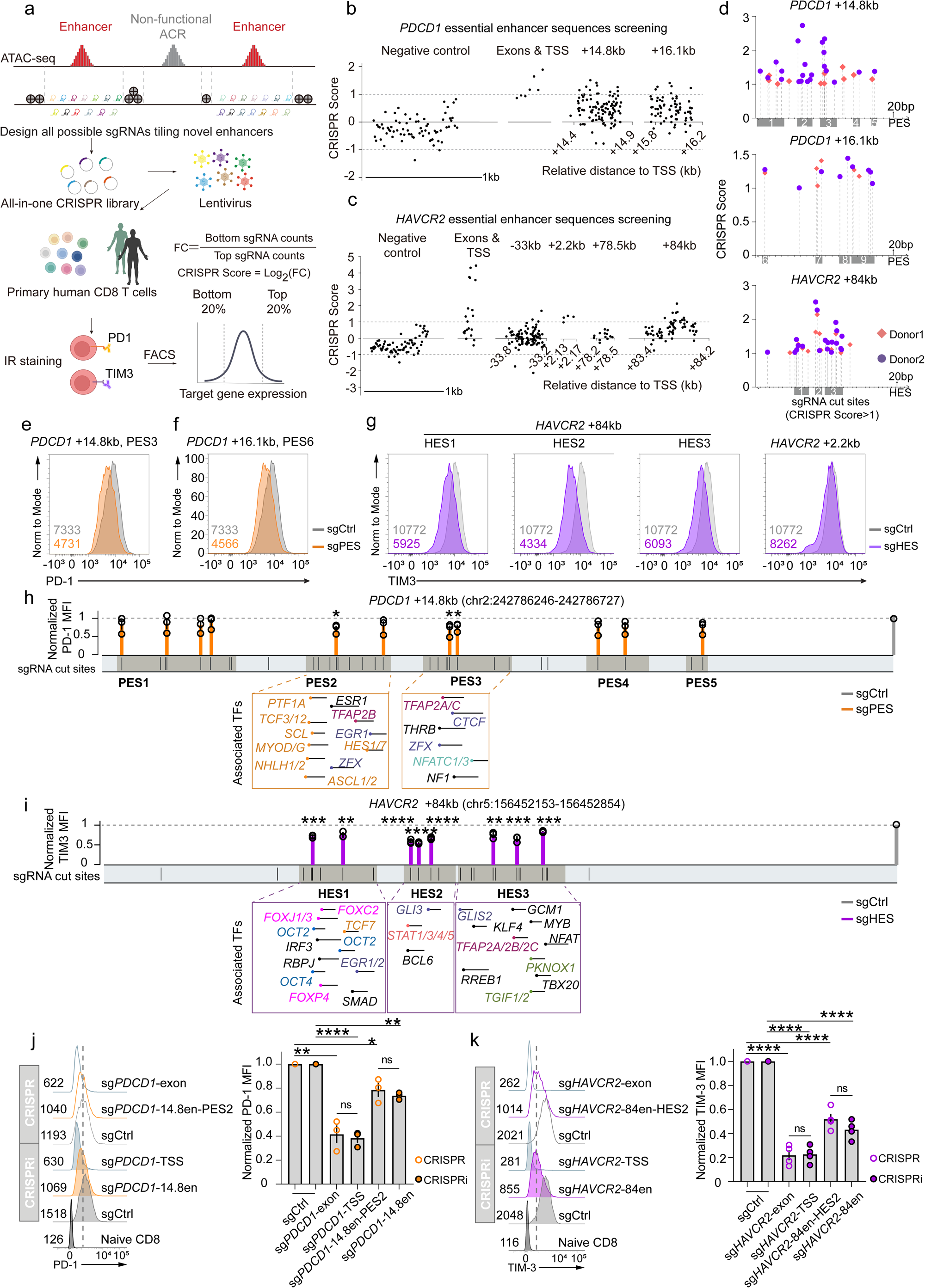
CRISPR saturation mutagenesis screening identifies essential components within *PDCD1* and *HAVCR2* CREs. **a**, Schematic of the CRISPR-based indel mutagenesis screens to map ESs within functional CREs of *PDCD1* or *HAVCR2*. **b,c,** ESs screening results for *PDCD1* (**b**) or *HAVCR2* (**c**) within the identified CREs (donor ND584). CRISPR score was calculated as the log2-fold change (Log_2_FC) in the abundance of the bottom versus the top bins. Each dot represents the CRISPR score and genomic distribution of an individual sgRNA. Negative control sgRNAs, as well as those targeting TSS and exons, were pseudo-mapped with a 1-bp interval. **d,** Genomic distribution of high-scoring sgRNAs (CRISPR score >1) within the *PDCD1* +14.8 kb, +16.1 kb, or *HAVCR2* +84 kb CREs. Shaded boxes indicate ESs with clustered sgRNAs. **e-g,** Representative flow cytometry plots showing Cas9-mediated disruption of PES3 (**e**), PES6 (**f**), and HES1-3 and +2.2kb CRE (**g**). Numbers on the bottom are the MFI of each curve, colors coded accordingly. **h,i**, Normalized protein expression of PD-1 (**h**) or TIM-3 (**i**) from validation of the identified *PDCD1* or *HAVCR2* ESs by Cas9-indel mutagenesis. Colored bars represent mean ± s.e.m. of 3 donors and three independent experiments. Black bars mark the predicted CRISPR DNA cleavage sites. Shaded areas indicate the ESs. Dashed boxes highlight the predicted TF binding motifs. **j,k,** Representative flow cytometry plots and summarized MFI of PD-1(**j**) or TIM-3(**k**) with Cas9-indel mediated disruption of individual ES or CRISPRi-mediated reduction of the entire CRE. Each dot represents a donor. Bar represents mean ± s.e.m. of 3-4 donors from two independent experiments. ns, not significant. *P<0.05, \*\**P* < 0.01, \*\*\**P* < 0.001 and ****P<0.0001, two-tailed Student’s t-test (h-k).

Fine-mapping using this Cas9-indel screening approach identified essential sequences (ESs) within the newly defined *PDCD1* and *HAVCR2* enhancers. Most negative control sgRNAs had CRISPR scores centered around 0, with background variation ranging from -1 to +1. sgRNAs with the highest CRISPR scores (positive regulators of the target gene) clustered around the coding sequences and promoter-proximal regions (Fig. 3b, c; Extended Data Fig. 7c, d). Although most indel-targeting sgRNAs within the CREs had CRISPR scores similar to negative controls, there were clusters of high-scoring sgRNAs at focal regions within the *PDCD1* +14.8 kb and +16.1 kb CREs, as well as in the *HAVCR2* +2.2 kb and +84 kb CREs (Fig. 3b, c and Extended Data Fig. 7c, d). Since Cas9-induced indels occur within ∼10 bp of the predicted DNA cleavage site, we used these data to define genomic sites covered by these sgRNAs with high CRISPR scores (Log2FC>1) as ESs within these CREs. Using these criteria, we identified five and four *PDCD1* ESs (PES1–5 and PES6–9) within the +14.8 kb and +16.1 kb CREs, respectively, and three *HAVCR2* essential sequences (HES1–3) within the +84 kb CRE (Fig. 3d). To extend these findings, we individually introduced Cas9-induced indels to CD8 T cells from a third donor using multiple high-scoring sgRNAs targeting the identified ESs. Flow cytometry confirmed on-target reduction in PD-1 and TIM-3 protein expression following disruption of the ESs within the *PDCD1* +14.8kb and +16.1kb CREs, as well as the *HAVCR2* +84 kb and +2.2kb CRE (Fig. 3e-g, Extended Data Fig. 7e, f). High scoring sgRNAs targeting PES2 and PES9 were initially from one donor in the screening data. We validated these sites using two donors, the consistent reduction of PD-1 expression in both donors confirmed the reproducibility of PES2 and PES9 (Extended Data Fig. 7f).

To evaluate the contribution of the short regulatory DNA sequences within ESs to overall gene regulation, we quantified the reduction in protein expression (MFI) after targeting the different ES regions. Disrupting PES2 and PES3 resulted in significant reduction in PD-1 expression compared to other PESs (Fig. 3h). Interfering with the HES1-3 within TIM-3 +84kb enhancer decreased TIM-3 expression, with HES2 having the strongest effect (Fig. 3i). Mapping the potential TF binding sites within PES2, PES3 and the three HESs using the MEME Suite revealed bHLH family motifs were enriched in PES2, whereas PES3 contained potential binding sites for TFAP2A/C, NFATC1/C3, and ZFX (Fig. 3h). For the *HAVCR2*, motifs from the STAT family, BCL6, and GLI3 were enriched most strongly in the HES2 site (Fig. 3i). To test the functional causality of these TF binding sites for the STAT family, we directly tested the role of *STAT5B* in regulating *HAVCR2* expression. Indeed, CRISPRi-mediated knockdown of *STAT5B* in CD8 T cells reduced TIM-3 expression by 27%, indicating a role for STATs in regulating expression of this IR (Extended Data Fig. 7g, h).

We next compared indel-based mutagenesis of a dominant ES with CRISPRi-mediated silencing of the corresponding full CRE to assess their relative effects on gene expression. For *PDCD1*, indel-based mutation of PES2 within the +14.8kb CRE resulted in similar reduction of PD-1 expression compared to full enhancer inhibition by CRISPRi (Fig. 3j). A similar result was observed at the *HAVCR2* +84kb enhancer, where targeting only the ES resulted in a ∼50-60% reduction in TIM-3 expression similar to targeting the entire CRE, with no significant difference between the two approaches (Fig. 3k). Together, these data define functional components within the IR CREs and establish PES2, PES3, and HES2 as critical elements governing the function of the *PDCD1* +14.8 kb and *HAVCR2* +84 kb enhancers. Targeted disruption of these individual elements was sufficient to largely abrogate the enhancer function in human CD8 T cells.

### Combinatorial enhancer perturbation identifies optimal enhancer pairs for fine-tuning inhibitor receptor expression

The results above identified multiple enhancers controlling *PDCD1* and *HAVCR2* expression, raising the question of whether these elements act in a coordinated manner to modulate IR expression. For the +14.8kb and +16.1kb CREs of *PDCD1* and +84kb CRE of *HAVCR2*, most sgRNAs had high CRISPRi scores, consistent with strong regulatory function. In addition to these major enhancers, the CRISPRi screens also identified additional regions for both IR genes with potential contributory enhancer activity. However, these ACRs often only had one or two sgRNAs that yielded high scores, suggesting more modest or context-dependent activity. These regions include *PDCD1* - 4.4 kb, -4.65 kb, and -38 kb ACRs, the *HAVCR2* +9 kb and +89 kb ACRs. High-scoring sgRNAs targeting these ACRs demonstrated clear PD-1 or TIM-3 expression reduction, confirming their enhancer function (Extended Data Fig. 8a, b).

To interrogate the potential combinatorial function of these enhancers in modulating *PDCD1* or *HAVCR2* expression, we performed multi-enhancer perturbation at the single-cell level. CD8 T cells were co-transduced with three all-in-one CRISPRi constructs, each targeting a distinct enhancer of *PDCD1* or *HAVCR2* and expression of PD-1 or TIM-3 was assessed (Fig. 4a, b). We targeted the *PDCD1* enhancers (PEN for the entire enhancer versus PES above for the specific sequence with the enhancer), PEN1 to PEN5 and *HAVCR2* enhancers (HEN), HEN1 to HEN5 as well as ACRs proximal to the TSSs as positive controls with sgRNAs using CRISPRi (Fig. 4c, g). Combinatorial perturbation of enhancers in individual CD8 T cells using CRISPRi led to additive reduction in expression of *PDCD1* or *HAVCR2* (Fig. 4d, h). At the *PDCD1* locus, three double-enhancer perturbations resulted in significantly stronger reduction of PD-1 expression compared to their respective single-enhancer perturbations, including: PEN2–PEN3 (+14.8kb─-4.4kb), PEN2–PEN4 (+14.8kb─-4.65kb) and PEN1–PEN3 (+16.1kb─- 4.4kb) (Fig. 4e, f and Extended Data Fig. 8c, d). For *HAVCR2*, seven specific double-enhancer combinations generated more effective regulation of TIM-3 expression compared to their corresponding single-enhancer perturbations, these enhancer-pairs are: HEN1-HEN2 (+89kb─+84kb), HEN2-HEN4 (+84kb─+9kb), HEN3-HEN5 (+78.5kb─-33kb), HEN1-HEN5 (+89kb─-33kb), HEN1-HEN4 (+89kb─+9kb), HEN1-HEN3 (+89kb─+78.5kb), and HEN3-HEN4 (+78.5kb─+9kb) (Fig. 4i-k and Extended Data Fig. 8e-i). Among these dual-enhancer pairs, co-targeting HEN2 and HEN4, resulted in the most robust reduction of TIM-3 expression (Fig. 4i). Co-targeting weak enhancer pairs also resulted in enhanced reduction of TIM-3 expression, but the magnitude varied for different combinations. Dual perturbation of HEN1–HEN3 produced a greater reduction in TIM-3 MFI compared to the HEN3–HEN5 and HEN1–HEN5 pairs, highlighting functional heterogeneity even among weak enhancer combinations. Notably, we identified two distinct triple-enhancer combinations: HEN1-HEN2-HEN4 (+89kb-+84kb-+9kb) and HEN1-HEN3-HEN4 (+89kb-+78.5kb-+9kb), that had regulatory effects comparable in magnitude to those observed when targeting the TSS (Fig. 4i, m). Indeed, targeting the HEN1-HEN3-HEN4 enhancer combination resulted in more robust control of *HAVCR2* expression than any of the corresponding double-enhancer combinations (Fig. 4i, l). Collectively, these data revealed the integrated regulation of *PDCD1* and *HAVCR2* by specific enhancer combinations, providing flexible strategies for modulating the expression of these IRs.

**Fig. 4:**
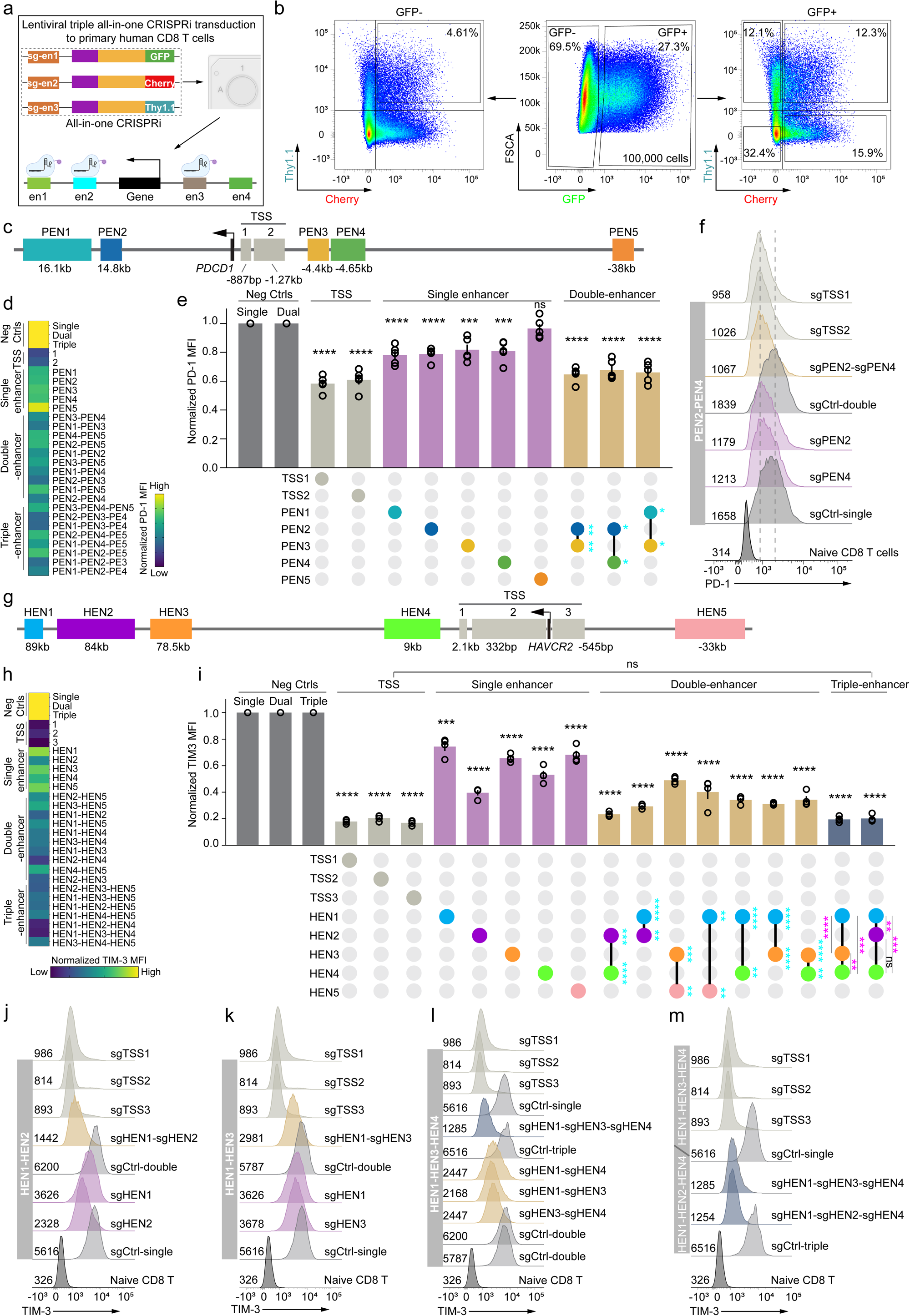
Combinatorial CRE perturbation identifies specific enhancer combinations for fine-tuning IR expression. a, Schematic of triple all-in-one CRIPSRi lentiviral transduction to primary human CD8 T cells. **b,** Representative flow cytometry plots showing the transduction efficiency of lentiviruses encoding three distinct all-in-one CRISPRi constructs. Cells gated as CD8+ live singlets with a minimum of 100,000 GFP+ cells were gated for further analyses. **c,** Genomic organization of *PDCD1* CREs identified in CRISPRi screens. **d**, Normalized PD-1 MFI across all single-, double-, and triple-CRE perturbation groups. PD-1 MFI values from single- and double-CRE perturbations were normalized to their respective single- and double-negative controls. **e,** Quantification of normalized PD-1 MFI of all single- and statistically significant double-*PDCD1* CRE perturbation groups. Bar represents mean ± s.e.m.; n = 4 donors from five independent experiments. Filled circles beneath histograms represent single- and double-CRE perturbations, the two perturbed CREs are connected by black lines. Targeted CREs are color-coded as in (**c**). Asterisks in black indicate significance relative to corresponding negative controls; asterisks in blue indicate significant improvement of dual- over single-CRE perturbations. **f,** Representative flow cytometry plots of PD-1 expression for the indicated CRE perturbations, numbers on bottom left represent MFI. **g**, Genomic organization of the identified *HAVCR2* CREs from CRISPRi screens. **h,** Normalized TIM-3 MFI across all single-, double-, and triple-CRE perturbation groups. TIM-3 MFI values from single-, double- and triple-CRE perturbations were normalized to their respective single-, double- and triple-negative controls. **i,** Quantification of normalized TIM-3 MFI for all single-and statistically significant double- and triple-*HAVCR2* CRE perturbation groups. Bar represents mean ± s.e.m. of 3 donors from four independent experiments. Filled circles under histograms denote single-, double-, and triple-CRE perturbations, with perturbed CREs connected by black lines; CREs color-coded as in (**g**). Asterisks in black indicate significance versus negative controls; blue, significance of double versus single perturbations; pink, triple versus double perturbations. ns, not significant. *P<0.05, **P < 0.01, ***P < 0.001 and ****P<0.0001, two-tailed Student’s t-test. **j-m,** Representative flow cytometry histograms of TIM-3 expression with indicated single-, double-, and triple-*HAVCR2* CRE perturbations, numbers on bottom left indicate MFIs of each curve.

### Dynamic use of *PDCD1* and *TIM3* enhancers during CAR T cell dysfunction

Expression of PD-1 and TIM-3 is transiently induced upon T cell activation. However, chronic antigen stimulation leads to T cell exhaustion, which is characterized by sustained co-expression of multiple IRs that inhibit optimal T cell function. This process is accompanied by extensive transcriptional and epigenetic reprogramming that remodels the chromatin landscape, including at IR gene loci. The *PDCD1* and *HAVCR2* enhancers defined above were identified following acute antigenic stimulation. To interrogate the extent to which these enhancers might be differentially active during prolonged T cell stimulation, we used an anti-GD2 CAR that induces persistent tonic signaling and recapitulates hallmark features of T cell exhaustion^38,39^. Long-term in vitro survival of chronically stimulated CD8 T cells can be augmented by including CD4 T cells^40^. Thus, CD8 T cells were co-transduced with an all-in-one CRISPRi construct targeting *PDCD1* or *HAVCR2* enhancers along with the anti-GD2 CAR; donor-matched CD4 T cells were transduced with the CAR alone. These CD4 and CD8 T cells were then co-cultured at a 1:1 ratio for up to 31 days, a setting that supports long-term survival of chronically stimulated CD8 T cells. CD8 T cell differentiation and expression of PD-1 and TIM-3 were assessed over time (Fig. 5a). As expected, this anti-GD2 in vitro co-culture model resulted in a marked decrease in CD62L^-^CD45RO^-^ (central memory, CM) and CD62L^+^CD45RO^-^(stem cell memory, SCM) CD8 T cells and corresponding increase in more effector-like CD62L^-^CD45RO^-^ (E) and CD62L^-^CD45RO^+^ (effector memory, EM) cells by day 31 (Extended Data Fig. 9a). These shifts in T cell subset distribution were consistent across all edited groups (Extended Data Fig. 9b, c).

**Fig. 5:**
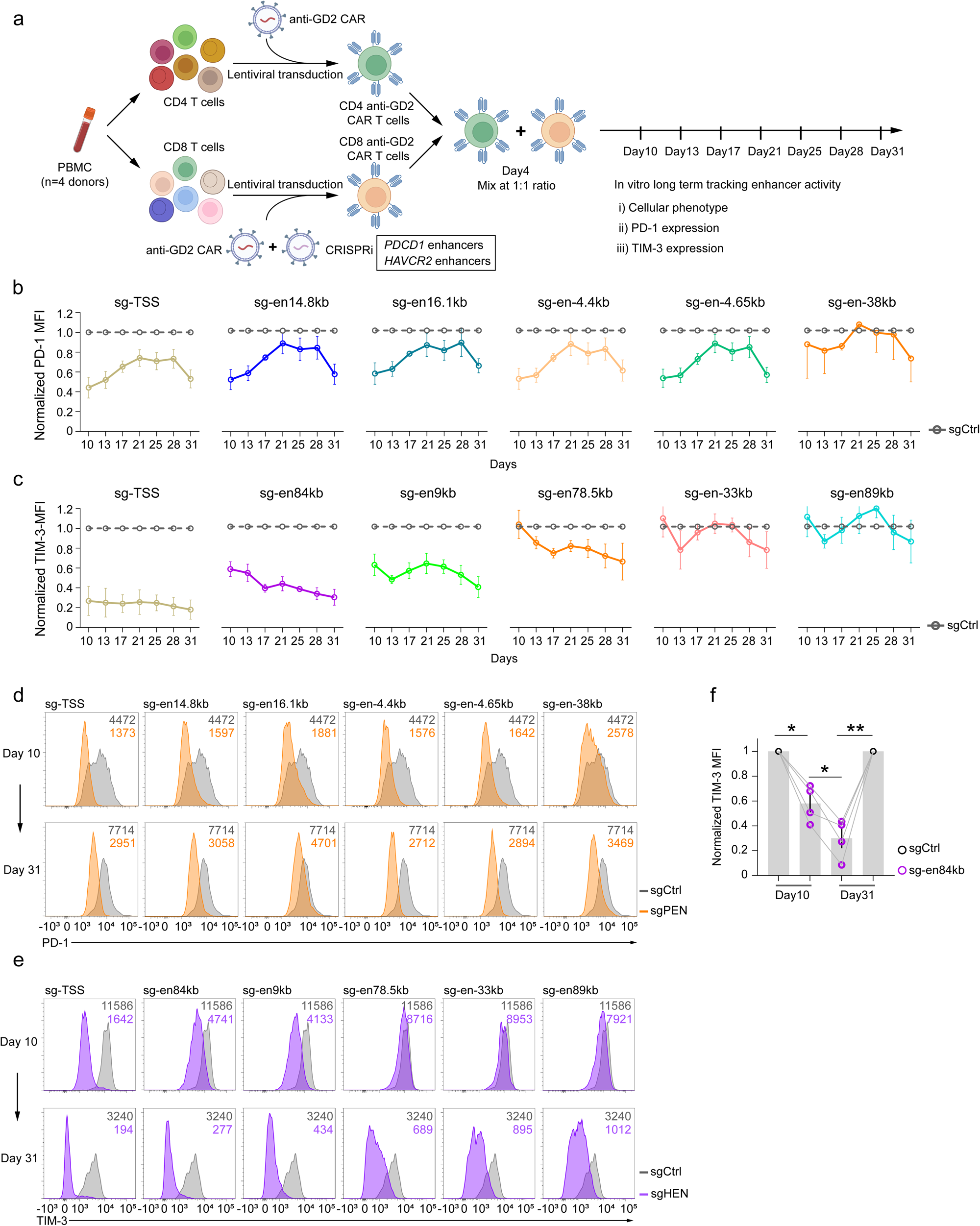
Dynamics and perturbation of IR CREs during chronic stimulation of CAR T cells. a, Schematic illustrating the tracking of CRE activity and effects of CRISPRi perturbation using an in vitro anti-GD2 tonic signaling CAR T cell model over time. **b,c,** Temporal dynamics of normalized PD-1 (**b**) or TIM-3 (**c**) expression with TSS or CREs targeted as indicated. Lines represent mean ± s.e.m. at each time point, n = 4 donors from four independent experiments. Lines are color-coded as shown in Fig. 4c, g. **d,e,** Representative flow cytometry histograms of PD-1 (**d**) or TIM-3 (**e**) expression at days 10 and 31 with TSS or CREs targeted by CRISPRi as indicated. MFI on plot as indicated, colors coded accordingly. **f,** Comparison of TIM-3 expression at days 10 and 31 following perturbation of *HAVCR2* +84kb CRE edited. Bar represents the mean ± s.e.m;, n= 4 donors form four independent experiments. Lines connect individual donors. P<0.05, **P < 0.01, two-tailed Student’s t-test.

Temporal analysis of PD-1 and TIM-3 expression in CAR+ and CRISPRi+ cells revealed dynamic patterns of expression. Targeting the *PDCD1* TSS initially resulted in efficient reduction of PD-1 expression, but this inhibitory effect gradually waned from day 10 to 28 with a recovery of suppression by day 31 (Fig. 5b). Similar dynamic patterns were observed for enhancer-targeted groups (Fig. 5b). All of the *PDCD1* enhancers had their strongest impact during the initial ∼3 weeks of culture (Fig. 5b, d; Extended Data Fig. 9d) when development of T cell exhaustion is being established.

The pattern of regulation of TIM-3 was distinct from PD-1. CRISPRi targeting the *HAVCR2* TSS induced robust and sustained reduction in gene expression (Fig. 5c). Among the enhancers, targeting the +84 kb and +9 kb elements resulted in early reduction of TIM-3 expression that became stronger over time (Fig. 5c), consistent with the gradual increase in TIM-3 gene expression as T cell exhaustion advances^41^. However, although the +84 kb and +9 kb enhancers initially exerted comparable effects on TIM-3 expression, the potency of these two enhancer elements diverged from day 13 onward. CRISPRi mediated targeting of the +84 kb enhancer reduced TIM-3 expression ∼40% on day 10 but this effect increased to ∼70% by day 31, nearly approaching the efficiency achieved by targeting the TSS (Fig. 5c). The progressive decline in TIM-3 expression (MFI) over time suggested that the activity of this enhancer increased steadily as T cell differentiation progressed to terminal exhaustion (Fig. 5f). Furthermore, from day 17 onward, the TIM-3 MFI in cells edited at the +84 kb enhancer remained lower than in cells subjected to other enhancer perturbations, underscoring the dominant role of this +84kb enhancer in regulating *HAVCR2* expression. In contrast, the +9 kb enhancer had a biphasic impact, with increased activity at day 13, declining modestly until day 21, then increasing again from day 25 to day 31 (Fig. 5c). The impact of the +78.5 kb enhancer was negligible at day 10 but became stronger as T cell differentiation progressed (Fig. 5c, 5e; Extended Data Fig. 9e). Intriguingly, the −33 kb and +89 kb enhancers showed no detectable activity before day 28, but at later time points these enhancers showed modest control over TIM-3 expression, implicating these regulatory regions in control of *HAVCR2* expression at later dysfunctional stages (Fig. 5c, e; Extended Data Fig. 9e). Together, these data reveal distinct dynamic patterns of enhancer use by *PDCD1* versus *HAVCR2* over time during T cell differentiation in this in vitro model. In particular, the *PDCD1* +14.8kb, 16.1kb, -4.4kb and -4.65kb enhancers, the *HAVCR2* +84kb, +78.5kb and +9kb enhancers, remain functionally active even at late stage of long-term in vitro induction.

### Fine-tuning inhibitory receptor expression via enhancer editing enhances CAR T cell efficacy

Targeting IRs through antibody blockade has demonstrated substantial therapeutic efficacy in cancer treatment, highlighting the potential of genetic engineering to harness these regulatory pathways. However, complete genetic loss of IRs can exacerbate exhaustion, impair long-term responses^23^, and prevent formation of proper T cell memory^9,24^. We therefore investigated how genetic targeting of the non-coding DNA elements controlling IR expression, rather than full IR KO, would compare for optimizing CAR T cell biology. Here, we focused on TIM-3. In vitro, manipulating TIM-3 function on human T cells using antibodies can improve effector function and lead to preservation of CD62L^+^ central memory (CM) like differentiation^42,43^. We used two approaches to modulate expression of this IR in anti-GD2 targeting CAR T cells. First, *HAVCR2* was genetically ablated by CRISPR-Cas9-mediated targeting of coding exons. Second, we targeted HES2 (within the +84kb enhancer) to fine-tune *HAVCR2* expression (Fig. 6a). Engineered anti-GD2 CAR T cells were cultured with 143B tumor cells that were replenished every four days to maintain the same CAR T cell to tumor ratio over time (Fig. 6a) and TIM-3 editing efficiency was consistent and persistent over time (Fig. 6b). On day 4, CAR T cells with a control edit (sgCtl), edit of the coding sequence to generate a TIM-3 KO (sgTIM-3-exon), or edit of the +84kb HES2 (sgTIM-3-84kb-en) to generate a partial knock down (KD) of TIM-3 all resulted in robust tumor killing (Fig. 6c). However, by day 8 of co-culture with tumor cells, the control and TIM-3 KO CAR T cells lost the ability to fully control tumor cells whereas the +84kb HES2 edited anti-GD2 CAR T cells still retained approximately 50% killing efficiency (Fig. 6c, d), suggesting that the ability to sustain T cell responses due to HES2 editing rather than full KO of TIM-3 may contribute to better tumor control. Moreover, at day 4, the number of +84kb HES2 edited CAR T cells showed a trend towards higher numbers compared to other groups (Fig. 6e). TIM-3 HES2 edited CAR T cells also produced more IFNγ and GM-CSF than full KO CAR T cells, though the enhancer edited and control edited cells were similar for these functions (Fig. 6f).

**Fig. 6:**
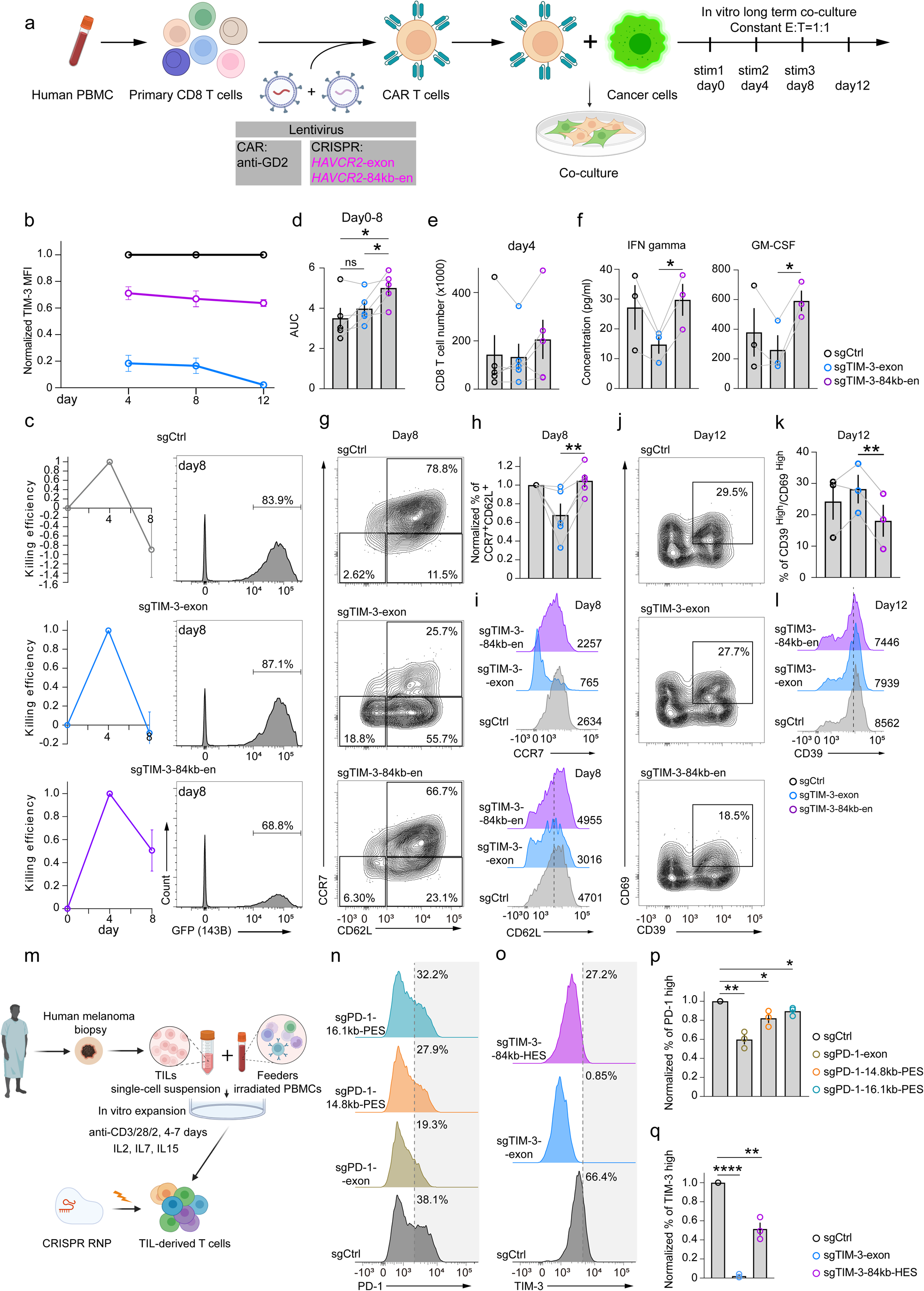
Fine-tuning *HAVCR2* expression via CRE editing enhances anti-GD2 CAR T fitness and modulates IR expression in melanoma TIL-derived CD8 T cells. **a**, Schematic experimental design of in vitro co-culture of IR-targeting genome-edited anti-GD2 CAR T cells with 143B osteosarcoma tumor cells. CAR T cells were subjected to either indel mutagenesis of an exon or targeted enhancer perturbation. Edited CAR T cells were analyzed on days 4, 8, and 12 of co-culture with GFP+ 143B osteosarcoma. **b,** Normalized expression of TIM-3 by CAR T cells at each time point (normalized to the negative control at each time point). **c,** Quantification of tumor cell killing efficiency in negative control, *HAVCR2* coding exon, or CRE edited anti-GD2 CAR T cells at each time point (left). Right panel. Representative flow cytometry plots showing the proportion of GFP+ 143B osteosarcoma cells on day 8. Cells were gated as live singlets. Killing efficiency was calculated by comparing the number of viable 143B cells after co-culture to the initial tumor cell count determined based on the E:T ratio. **d,** Quantification of tumor killing efficiency in negative control, *HAVCR2* coding exon, or CRE edited anti-GD2 CAR T cells from day 0 - day 8 using area under the curve (AUC; area above 0) calculated from (**c**). Lines connect individual donors. **e**, Total CAR T cell numbers on day 4. **f,** Cytokine concentration measured from culture supernatant on day 8. Bars represent mean ± s.e.m. of 3 donors. Lines connect individual donors. **g,** Representative flow cytometry plots of T cell subsets as assessed by CD62L and CCR7 expression in control, *HAVCR2* exon, or CRE edited anti-GD2 CAR T cells on day 8. **h,** Proportion of CD62L^+^CCR7^+^ cells from (g) across donors. **b-e**, **h**, Data represents mean ± s.e.m.; n = 5 donors from three independent experiments, Lines connect individual donors. **i,** Representative flow cytometry histograms showing the expression of CD62L and CCR7 by anti-GD2 CAR T cells with *HAVCR2* exon or CRE editing as indicated on day 8. MFI values are shown at the bottom right. **j,** Flow cytometry analysis of T cell subsets based on CD39 and CD69 expression in control, *HAVCR2* exon and CRE edited anti-GD2 CAR T cells on day 12. **k,** Summary of CD39^High^CD69^High^ subsets from (**j**). Bars represent mean ± s.e.m.; from 3 donors. Lines connect individual donors. **l,** Representative flow cytometry plots for CD39 expression in anti-GD2 CAR T cells with *HAVCR2* exon or CRE edited as indicated, numbers on the bottom right denote MFI. **m**, Schematic of modulating IR expression in melanoma TIL-derived CD8 T cells by CRISPR-mediated CRE-editing. **n,o,** Representative flow cytometry histograms of PD-1 (**n**) or TIM-3 (**o**) expression in CD8 T cells from expanded TIL with coding exon or CREs editing as indicated. Gray shaded areas indicate the PD-1^High^ or TIM-3^High^ populations, numbers denote proportions. Cells were gated as live CD8+ singlets. **p, q,** Summary of normalized PD-1^High^ frequency (**p**) from (n) and TIM-3^High^ frequency (**q**) from (o). Bars represent mean ± s.e.m. of 3 donors from three independent experiments. *P<0.05, **P < 0.01, ***P < 0.001 and ****P<0.0001, two-tailed Student’s t-test.

We next investigated whether altering the expression of TIM-3 via HES2 versus exon editing impacted the differentiation patterns of CAR T cells. Complete KO of TIM-3 resulted in substantial loss of CCR7 expression and accumulation of CD62L^Low^ CCR7^Low^ cells. Partial TIM-3 reduction via targeting HES2 within the +84kb enhancer in anti-GD2 CAR T cells preserved a substantial proportion of CD62L^+^CCR7^+^ cells, comparable to the control sgRNA group (Fig. 6g-i). Targeting HES2 rather than a TIM-3 coding exon also resulted in lower CD39 and CD69 expression, consistent with less exhaustion or terminal differentiation (Fig. 6j-l; Extended Data Fig. 10a). Consistent with these observations, editing of HES2 in the +84 kb enhancer resulted in fewer CD69^High^TCF1^Low^ anti-GD2 CAR T cells at day 12 (Extended Data Fig. 10b). Together, these findings highlight the utility of targeting enhancers to fine tune the expression of key genes in T cells. In particular, such state-specific regulatory elements might provide opportunities to tune the function of genes such as IRs where complete loss may be detrimental to optimal T cell biology.

Cellular therapies beyond CAR T cells, including tumor-infiltrating lymphocytes (TIL), may also benefit from engineering non-coding regions to improve anti-tumor activity. Optimization of TIL therapy has gained particular clinical relevance since FDA approval of TIL therapy was recently granted^44,45^. We therefore tested whether editing *PDCD1* and *HAVCR2* CREs could modulate IR protein expression in human melanoma TILs. First, analyses of published ATAC-seq data confirmed the presence of the +14.8kb and +16.1kb ACRs of the *PDCD1* locus, as well as the *HAVCR2* +84kb ACR, in CD8 TILs from human melanoma (Extended Data Fig. 10c).

Next, we tested whether targeting these CREs resulted in decreased PD-1 or TIM-3 expression in CD8 T cells from melanoma TIL (Fig. 6m). First, TILs were expanded in vitro with irradiated PBMCs as feeder cells and high dose IL-2. CD8 T cells from the expanded TIL exhibited high expression of multiple key exhaustion-associated molecules, including PD-1, TIM-3, LAG-3, CD39, CD94, CTLA-4 and CD95 compared to CD8 T cells from TIL biopsy (Extended Data Fig. 10d). The proportions of PD-1^+^TIM-3^+^, as well as PD-1^+^CD39^+^ and PD-1^+^CTLA-4^+^ subsets were increased post expansion consistent with expansion of tumor-specific populations (Extended Data Fig. 10e). These data indicate that expanded CD8 TILs recapitulate key features of exhausted T cells, providing a source of cells to test manipulation of IR enhancers in clinically relevant exhausted CD8 T cells.

Electroporation of Cas9-sgRNA ribonucleoprotein (RNP) complexes targeting the ESs within *PDCD1* enhancers, PES2 (+14.8 kb) or PES9 (+16.1 kb), significantly reduced the proportion of PD-1^High^ TIL-derived CD8 T cells (Fig. 6n, p). Targeting HES2 within *HAVCR2*+84 kb enhancer led to significant reduction in TIM-3 expression in the expanded TIL-derived CD8 T cells (Fig. 6o, q). Notably, targeting PES2 in the +14.8 kb *PDCD1* enhancer resulted in an 18% reduction in PD-1 expression, similar to what was observed in CD8 T cells from healthy human PBMCs shown in previous data (Extended Data Fig. 10f). Similarly, targeting the HES2 decreased TIM-3 expression by 41%, comparable to the effects observed in PBMC-derived CD8 T cells (Extended Data Fig. 10g). These findings demonstrate that the enhancers identified through the CRISPRi screen outlined above provide actionable targets for fine-tuning IR expression in human TILs and more broadly support the strategy of engineering CREs in therapeutically relevant T cell populations.

## Discussion

Cell therapies are revolutionizing the treatment of hematologic malignancies and beginning to show promise beyond leukemias and lymphomas^46,47^. Genome engineering has received considerable attention as a strategy to improve cell therapies, with the potential to improve CAR T cells or TILs for the treatment of solid tumors or other diseases. In particular, CRISPR-based genome engineering and other approaches now enable targeted knock out of negative regulators or insertion of synthetic genes to enhance T cell function offering major potential advances for adoptive therapies^48,49^. Although there have been some improvements, engineered adoptive T cell therapies often fail due to T cell exhaustion, dysfunction, or suboptimal differentiation^50,51^. These T cell differentiation trajectories are now recognized to involve extensive epigenetic remodeling, including the use of state-specific CREs or enhancers that drive transcriptional programs such as exhaustion-specific gene expression^4,5^. In mice, for example, an enhancer at -23.8kb in the *Pdcd1* locus controls expression of PD-1 exclusively in exhausted CD8 T cells^4,27^. Deletion of this enhancer improves differentiation and preserves function^28^, whereas full deletion of PD-1, although initially beneficial, ultimately results in long-term deficiencies^9, 23,24,48^. Thus, targeting non-coding enhancer elements to achieve differentiation state-biased regulation of gene expression has the potential to provide new opportunities to control gene expression and ultimately function for cell therapy. It has been challenging, however, to comprehensively interrogate the causality and function of non-coding elements in human T cells because of limitations in the functional annotation and technical barriers to efficiently engineer the relevant enhancers of key genes of interest.

In this study, we established a framework for systematic interrogation of non-coding DNA elements in primary human T cells and assessed the function of these enhancers in regulating gene expression and T cell differentiation. First, we developed a high-efficiency CRISPRi platform that enables scalable interrogation of non-coding DNA elements in primary human T cells. Second, using this platform, we identified multiple functional enhancers controlling critical immune regulators, including *PDCD1, HAVCR2,* and *TBX21*, showing that these regulatory elements act both independently and cooperatively to tune gene expression. Third, combinatorial targeting of enhancers enabled graded rather than binary control of immune checkpoint expression, offering a level of precision not achievable through complete gene knockout. Finally, targeting newly discovered IR enhancers in melanoma TIL and CAR T cells reduced surface IR expression. Moreover, fine-tuning *HAVCR2* expression via enhancer editing in anti-GD2 CAR T cells enhanced a favorable differentiation state, reduced exhaustion, and increased effector cytokine production upon tumor coculture, compared to complete *HAVCR2* knockout. Together, these findings highlight a strategy for precision engineering of immune checkpoint expression via enhancer targeting, enabling rational design of next generation cell therapies.

Enhancer activity is dynamic across the genome and undergoes major remodeling during T cell differentiation and transitions between different T cell states or subsets (e.g. naïve T cells → effector T cells; stem-like precursors → memory or exhausted T cells). Exhausted T cells possess a distinct epigenetic landscape compared to more functional T cell subsets. Indeed, exhaustion-specific enhancers and epigenetic “scars” in exhausted CD8 T cells can limit the function and responses to reinfection or immunotherapy^52–55^. Overcoming the epigenetic constraints associated with T cell exhaustion or dysfunction remains a key goal for restoring optimal T cell responses. However, genome-wide functional enhancer mapping in low abundance cell populations, such as antigen-specific exhausted T cells, remains challenging. Here, we used activated human primary CD8 T cells to perform large genomic region CRISPRi screens to discover CREs, followed by Cas9-induced indel mutagenesis screens to identify essential sequences within key enhancer regions. Then, to test how these regulatory elements controlled gene expression in exhausted T cells, we leveraged in vitro exhaustion assays including repetitive stimulation models and tonic signaling CAR T cells, that approximate some aspects of in vivo human T cell exhaustion. Finally, we tested the function of the newly discovered PD-1 and TIM-3 enhancers in CD8 T cells from human melanoma. Thus, our all-in-one CRISPRi platform enabled large-scale genomic region multi-layered screens to discover functional CREs that regulate key genes in exhausted CD8 T cells. Moreover, CRISPRa-mediated enhancer activation and CRE-driven reporter assays highlight how this CRISPR-based platform can be used for both disruption and induction of state-specific gene regulation, and even potentially to enable construction of synthetic regulatory circuits. The approach outlined here can easily be applied to other genes and scaled to integrate other single-cell approaches to generate high-resolution, cost-effective maps of enhancer function^56,57^. Such strategies would enable systematic definition of the epigenetic regulatory circuits that shape T cell fate and function.

IRs are key regulators of T cell activation and function, and have a prominent role in T cell exhaustion^58^. Indeed, blocking the PD-1 pathway has revolutionized cancer therapy through reinvigorating exhausted CD8 T cells^59,60^ and targeting the PD-1 pathway or TIM-3 can reverse exhaustion in preclinical models^41,61^. Although work continues on developing TIM-3-based targeting approaches for human T cell therapeutics^62^, five FDA approved therapies targeting the PD-1/PD-L1 pathway highlight the clinical relevance of these targets. However, complete loss of PD-1 signals can cause defects in T cell memory and erode the durability of long term of antigen-specific CD8 T cell responses in chronic infections and cancer^9,23,24^. Strategies that reduce and/or modulate IR expression in a more regulated manner, but do not completely ablate signals from these pathways may strike a balance between limiting negative regulation and preserving T cell fitness. Indeed, targeting an enhancer upstream of the *Pdcd1* locus in mice decreases exhaustion^28^ but such approaches have not been investigated in human T cells. Using the human CRISPRi enhancer screening platform described here, we defined the functional CREs controlling *PDCD1* and *HAVCR2* expression in primary human CD8 T cells. We identified individual and combinations of enhancers that fine-tuned expression of these IRs and revealed a functional benefit for enhancer editing for regulating *HAVCR2* expression compared to complete loss of TIM-3 expression in CAR T cells. These results highlight the potential advantage of targeting enhancer elements rather than gene coding sequences to optimize T cell function in engineered T cells. Moreover, we identified enhancer-combinations capable of a wider range of expression control, offering more tunable strategies for controlling expression of PD-1 or TIM-3 to optimize T cell function. Given the recent FDA approval of TIL therapy^48,49^, the ability to apply our CRISPR enhancer engineering system to human TILs to target regulatory elements of *PDCD1* and *HAVCR2* extends this approach to clinically relevant cells. In the future, enhancer editing of tumor-specific cells may offer a strategy to optimize TIL-based cell therapies.

Beyond IRs, other genes represent promising targets for cell engineering. Nodal regulators such as TFs often coordinate suites of genes with related biological functions. TBET, for example, regulates genes involved in effector CD8 T cell biology, including homing, migration, and effector function^63^. However, high and sustained TBET expression can drive terminal differentiation and/or contribute to more severe exhaustion in some settings^64,65,66,67,68^. Thus, identifying the regulatory logic controlling *TBX21* expression in primary human CD8 T cells may enable selective augmentation of TBET expression in a more physiological and state-specific manner than enforced overexpression. By activating the *TBX21* enhancers with CRISPRa, we were able to not only increase *TBX21* expression but also enhance downstream cytotoxic genes such as GZMB and PRF1 (Perforin). These findings highlight the potential opportunities of targeting nodal regulators to rewire transcriptional networks in human T cells. It may be advantageous to tune *TBX21* expression by using natural non-coding enhancer elements, providing more subtle, but also cell state-specific regulatory control. This strategy could be expanded to other master regulators to provide an avenue for the design of next-generation CAR T and other engineered T cell therapies.

Thus, we demonstrate the potential to regulate human T cell biology by first functionally annotating and then engineering the non-coding genome. The all-in-one CRISPRi approach described here provides a robust platform and can be expanded to study the functional enhancers of other genes at even larger genomic regions. Moreover, there may be additional advantages to build upon this system in the future. Although CRISPR gene-edited cells have been used in humans for cell therapy trials, large DNA edits or deletions in the human genome has inherent risks due to off-target effects and potential chromosomal aberrations especially in settings of multiple edits^69^. These risks can be mitigated by employing non-DNA-damaging editing strategies, like base and prime editors, as well as other epigenome silencers^70–73^ to manipulating non-coding elements and control gene expression. Thus, it is now possible to functionally decode the non-coding gene regulatory networks of human T cells and use these annotated enhancer landscape maps to improve T cell biology with potential therapeutic opportunities for engineered cell therapies. These studies lay the foundation for next-generation, epigenetically engineered cell therapies.

## Acknowledgements

We thank all members of the Wherry lab and Shi lab, especially J.E.Wu for manuscript discussion and editing. We thank M.McLaughlin and C.Holliday for laboratory support. We also thank Y.Zhu, H.Sun and E.L.Park from Penn Human Immunology Core (P30 CA016520/ SCR_022380). This work was supported by NIH grants AI155577, AI115712, AI117950, AI108545, AI082630, AI149680, HL145754 (to E.J.W.), Parker Institute for Cancer Immunotherapy (to E.J.W.). Work in the Wherry lab is supported by the Parker Institute for Cancer Immunotherapy. J.S. is supported by NIH/National Cancer Institute (NCI) (R01 CA258904) and Parker Institute for Cancer Immunotherapy. P.W. is supported by a Momentum Fellowship from The Mark Foundation for Cancer Research. A.C.H. is supported by R01 CA273018, P50 CA261608, Damon Runyon Clinical Investigator Award, Pew-Stewart Stewart Scholars Program in Cancer Research. Huang lab is supported by Tara Miller Foundation and the Parker Institute for Cancer Immunotherapy.

## Contributions

P.W., J.R.G., J.S. and E.J.W. conceived and designed experiments. P.W. performed all experiments in this study. P.W. and J.R.G. performed flow cytometry sorting. J.S. designed the CRISPRi screening library. ATAC-seq data was analyzed by S.M. and J.R.G. P.W. and H.H. analyzed the CRISPRi/CRISPR screening data. A.C.H. provided frozen TIL specimens. P.W., J.R.G., J.S. and E.J.W. wrote the manuscript with the help from M.S.J.

## Extended Data Figures

**Extended Data Fig. 1:**
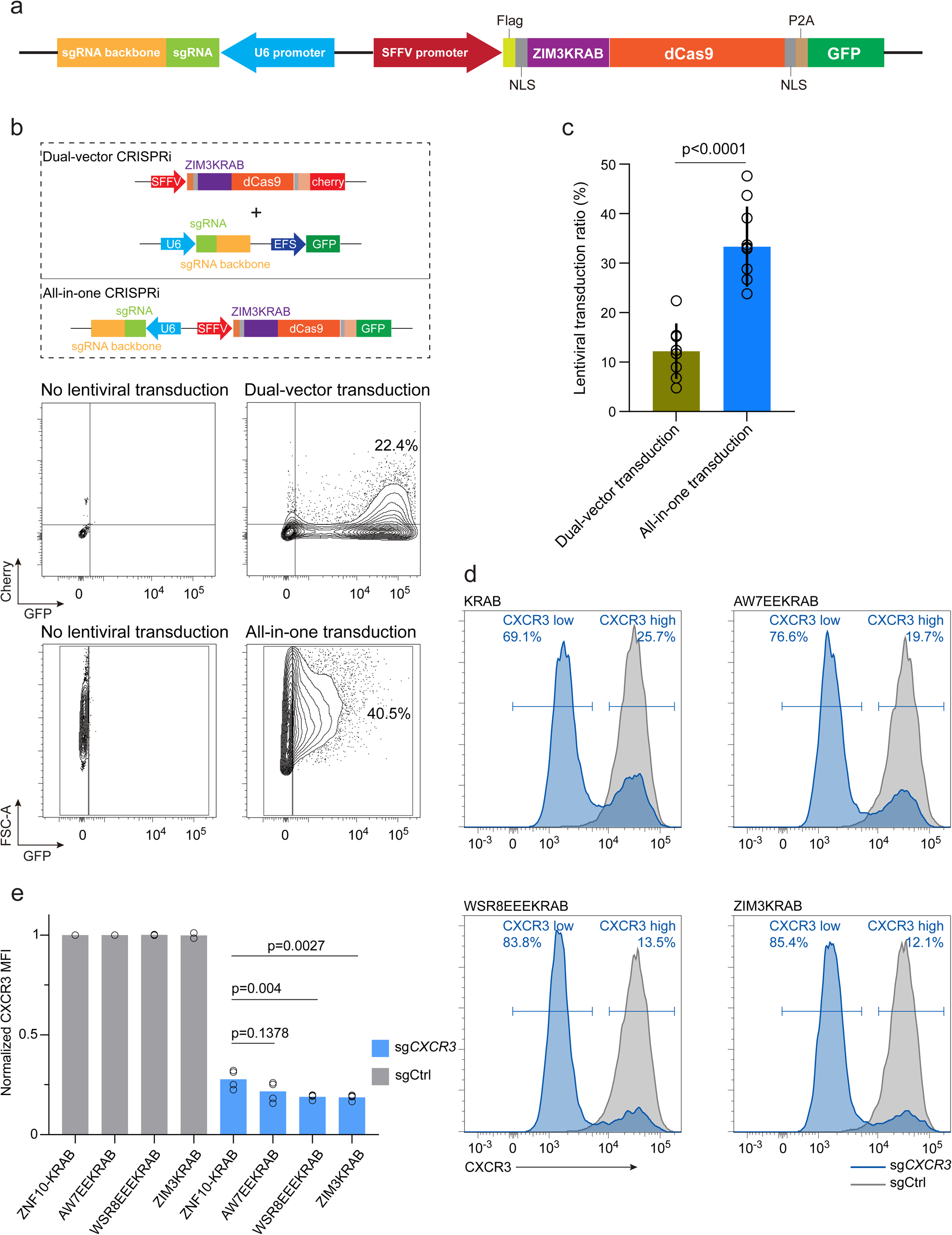
Development of all-in-one CRISPRi. **a**, Schematic of the all-in-one CRISPRi lentiviral vector design. **b**, Comparison of lentiviral transduction efficiency in primary human CD8 T cells between dual-vector CRISPRi and all-in-one CRISPRi. **c**, Quantification of lentiviral transduction efficiency using dual-vector CRISPRi and all-in-one CRISPRi (n=10 donors, mean±s.d., two-tailed Student’s t-test). **d**, Representative flow cytometry plots of CRISPRi-mediated CXCR3 reduction using all-in-one constructs incorporating different KRABs. **e**, Quantification of efficiency of reduction in CXCR3 expression mediated by KRAB variants (mean± s.d, n= 2 donors each with two technical replicates; two independent experiments, Two-tailed Student’s t-test).

**Extended Data Fig. 2:**
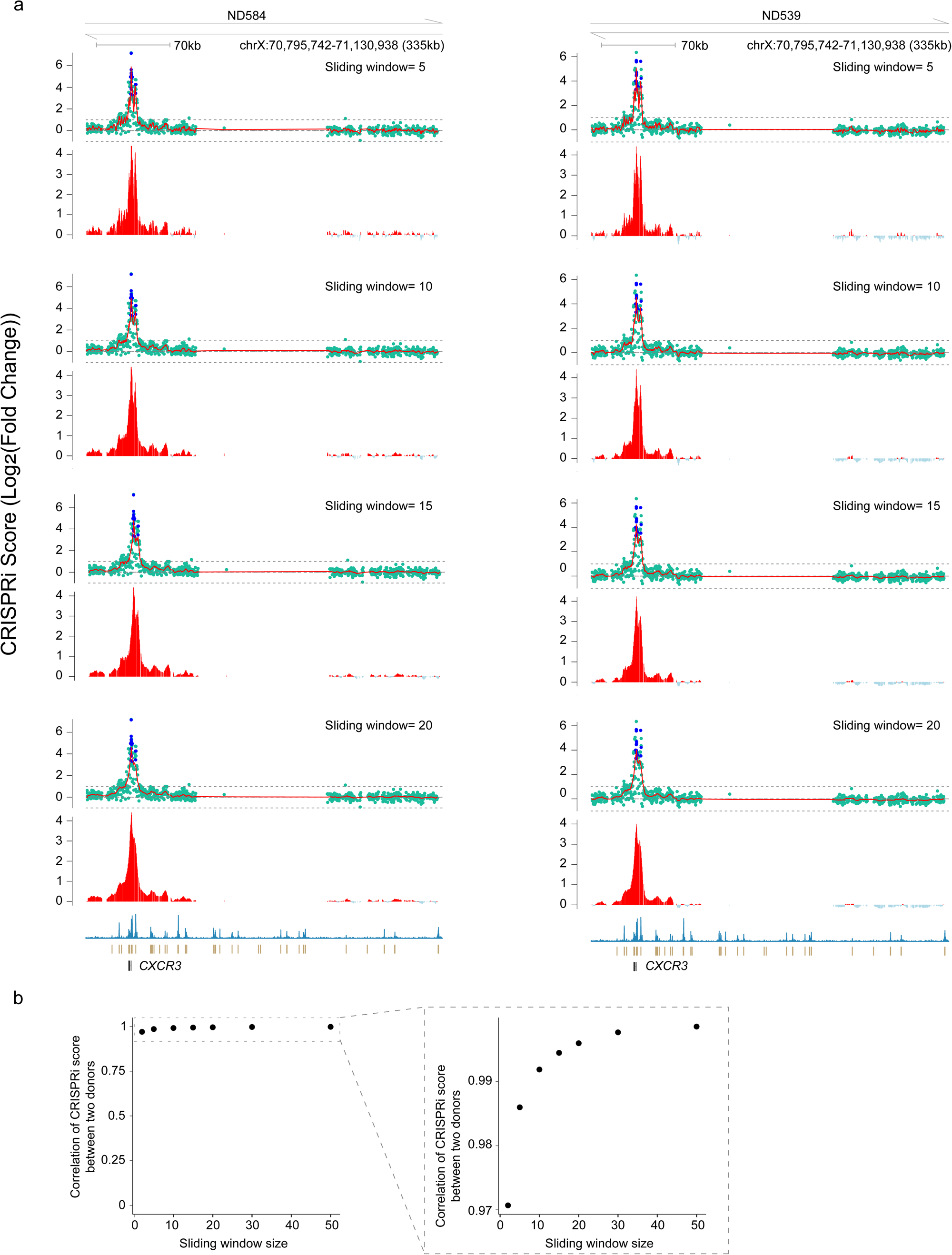
Deciphering the sliding window for tiled library screening. **a**, Tiled library pooled screening results for *CXCR3* after using different sliding window sizes in donor ND584 (left) and donor ND539 (right). Blue dots, positive controls. Green dots, designed sgRNAs from the tiled library. Red bars indicate CRISPRi scores calculated with the respective sliding window size, with red curves representing the same data as a smoothed line. **b**, Correlation of CRISPRi scores between the two donors across varying sliding window sizes. Dashed boxes highlight zoomed-in regions of the correlation plots.

**Extended Data Fig. 3:**
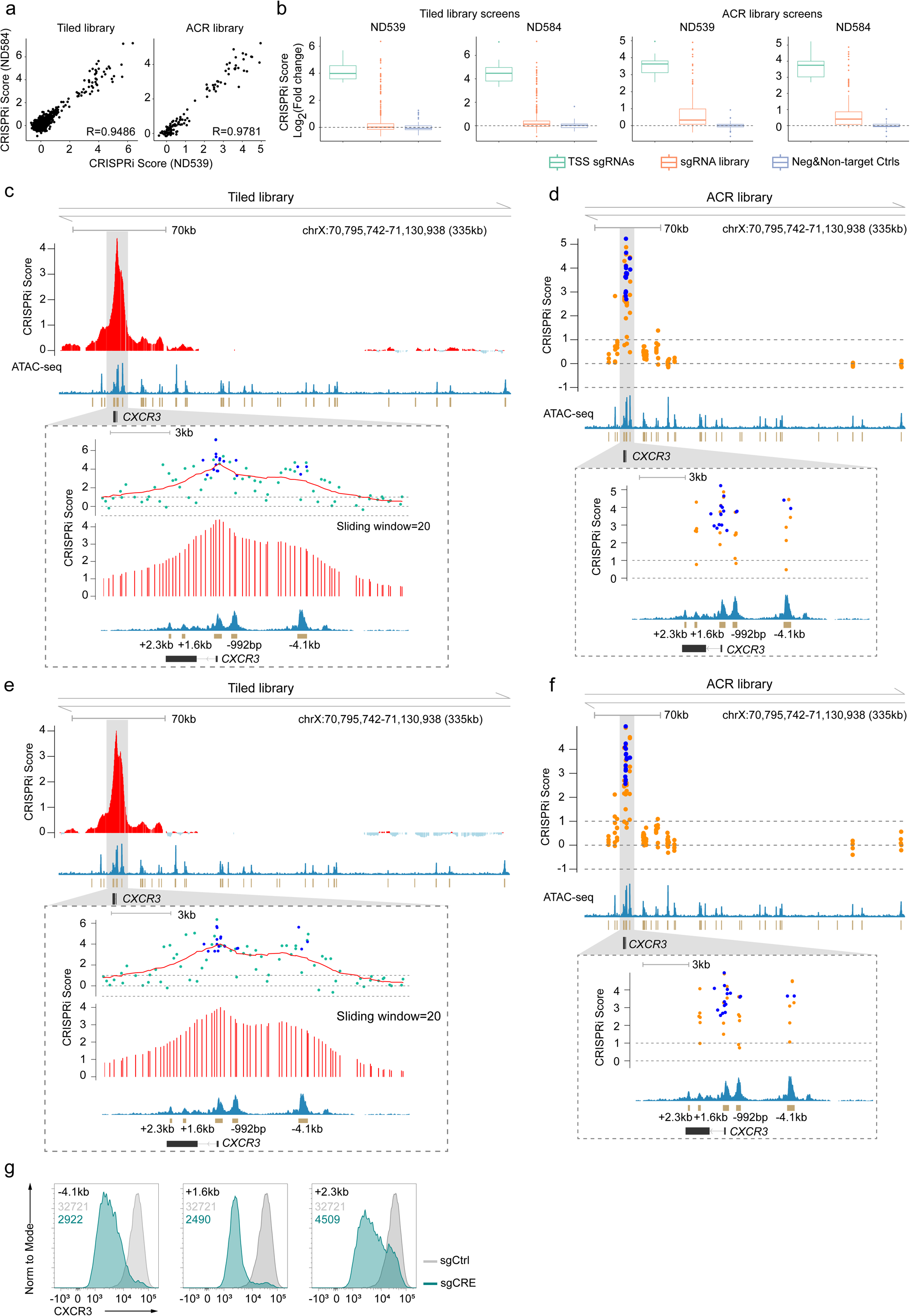
All-in-one CRISPRi screening for functional CREs of *CXCR3*. a, Scatter plot showing reproducibility of CRISPRi screening across two donors using the tiled library and ACR library. **b**, CRISPRi score distribution of each sgRNA across two donors in tiled and ACR screens. **c-f**, Tiled and ACR library screening results in donor ND584 (**c,d**) and donor ND539 (**e,f**). Red bars in c,e indicate CRISPRi scores calculated with a sliding window of 20; red curves show smoothed representations of the same data. Bottom, ATAC-seq profiles of the corresponding genomic regions. Gray shaded areas, zoomed-in regions with high CRISPRi score sgRNAs (Log2FC>1). Green dots, designed sgRNAs from tiled library. Blue dots, positive controls. Orange dots, designed sgRNAs targeting ACRs. **g**, Representative flow cytometry plots of CXCR3 expression following CRE targeting by CRISPRi. (Numbers on the top left indicate MFI, color coding matches the corresponding curves).

**Extended Data Fig. 4:**
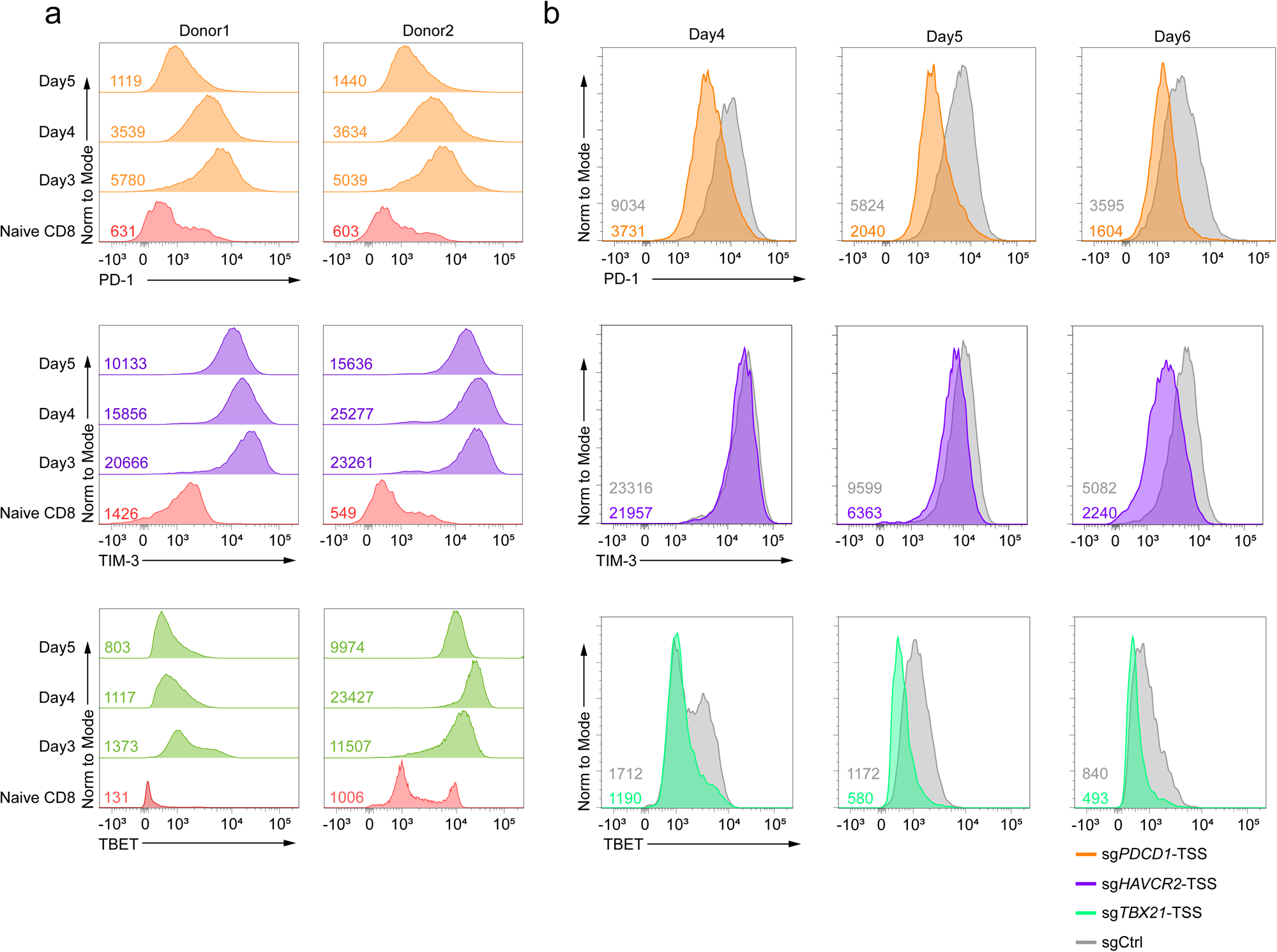
Expression dynamics and determination of optimal screening time points for *PDCD1*, *HAVCR2*, and *TBX21*. **a**, Flow cytometry plots of the expression patterns of PD-1 (top), TIM-3 (middle) and TBET (bottom) by in vitro cultured primary human CD8 T cells from two donors at different time points post-stimulation with antiCD3/28 beads. Red curve represents naïve CD8 T cells from human PBMC. **b**, Temporal reduction of PD-1, TIM-3, and TBET by CRISPRi in activated primary human CD8 T cells across corresponding time points. Numbers on the bottom left indicate the MFI of each curve, color coding matches the corresponding curves.

**Extended Data Fig. 5:**
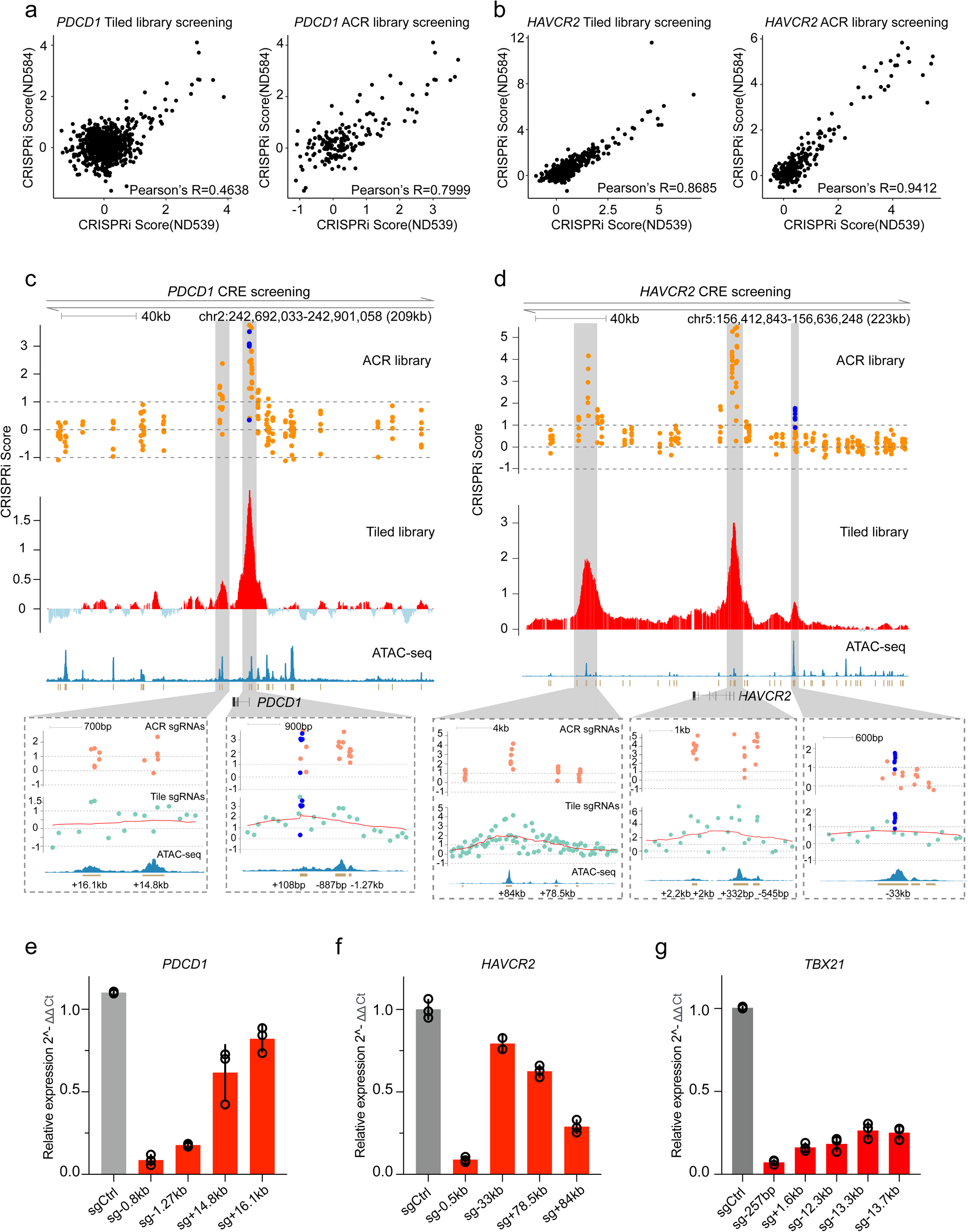
**All-in-one CRISPRi screening identified functional CREs of *PDCD1* and *HAVCR2*. a,b**, Scatter plot showing reproducibility of CRISPRi screening between two donors using both tiled and ACR libraries for *PDCD1* (**a**) or *HAVCR2* (**b**). **c,d**, Pooled CRISPRi screening results for functional CREs of *PDCD1* (**c**) or *HAVCR2* (**d**) using ACR and tiled library designs in donor ND539. Top, ACR library screening. Blue dots, positive controls. Orange dots, designed sgRNAs targeting ACRs. Middle, tiled library screening. Red bars, CRISPRi scores calculated with a sliding window of 20. Bottom, ATAC-seq profiles of the corresponding genomic regions showing chromatin accessibility. Gray shaded areas, zoom-in of regions with high CRISPRi score sgRNAs (Log2FC>1). Green dots, designed sgRNAs from the tiled library. Red curves represent the same data of red bars as a smoothed line. **e-g** Quantification of *PDCD1* (**e**), *HAVCR2* (**f**) or *TBX21* (**g**) mRNA expression by RT–qPCR after targeting the identified CRE by CRISPRi. Data represent mean ± s.d. from three technical replicates.

**Extended Data Fig. 6:**
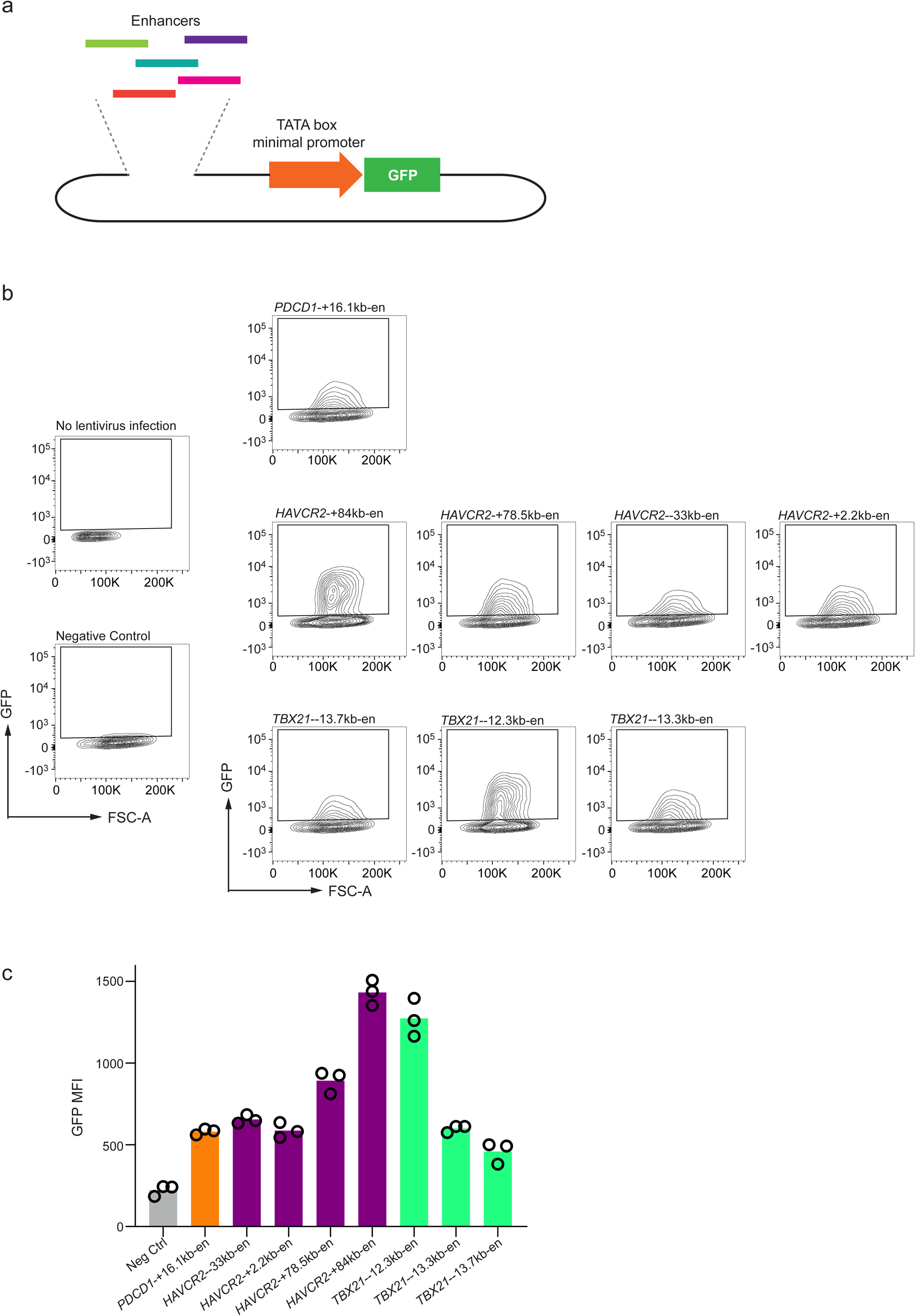
Reporter assay confirms CRE-mediated activation of target gene. **a**, Schematic of the reporter assay construct. **b**, Representative flow cytometry plots of GFP expression in primary human CD8 T cells after transduced with reporter constructs with different CREs. **c**, Histograms showing normalized GFP expression. Data represent mean ± s.d. from three technical replicates.

**Extended Data Fig. 7:**
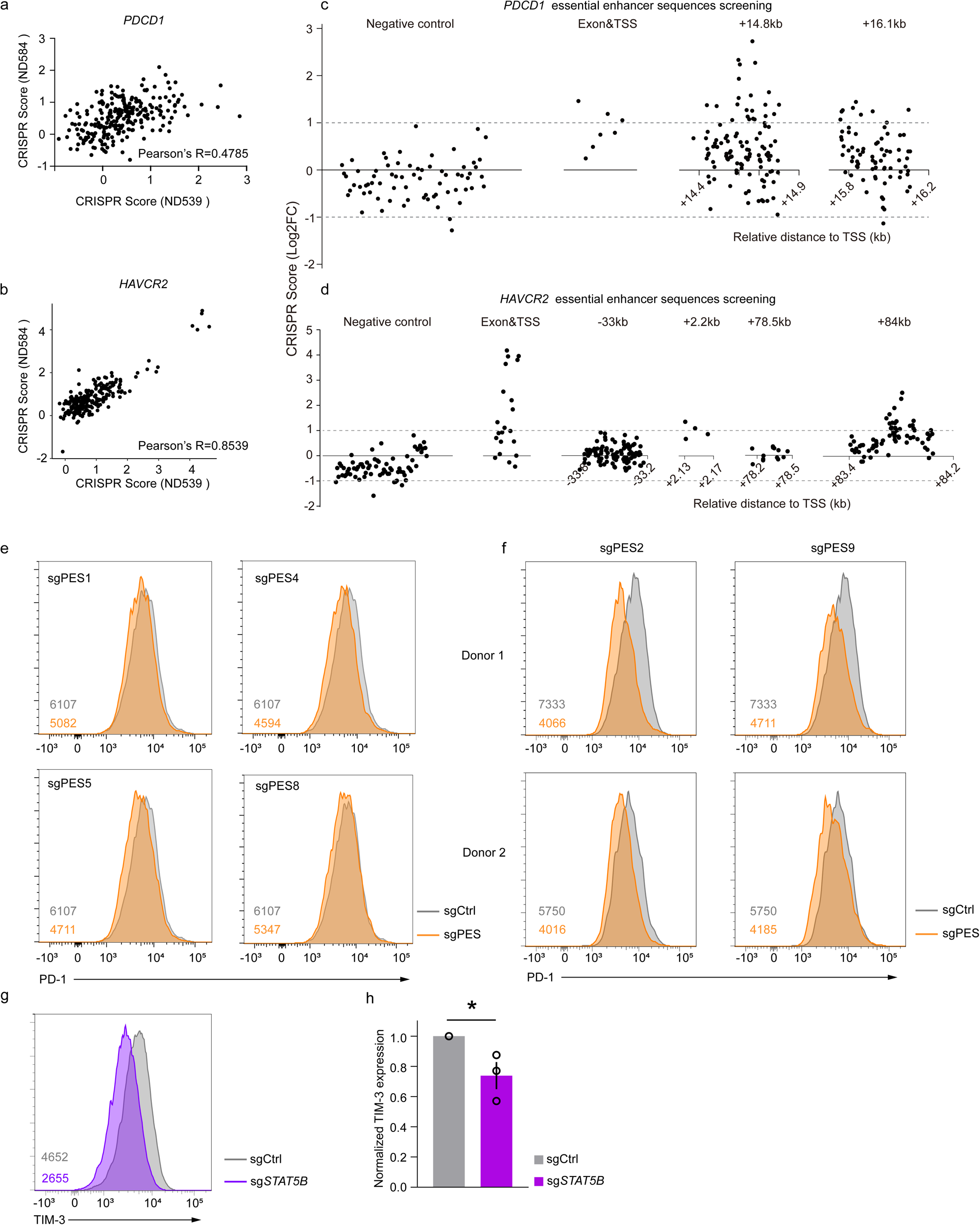
**CRISPR saturation mutagenesis screening revealed critical regulatory elements within *PDCD1* and *HAVCR2* CREs. a,b**, Reproducibility of *PDCD1* (**a**) and *HAVCR2* (**b**) screens across two donors. Pearson’s correlation. **c,d**, CRISPR screen results for *PDCD1* (**c**) and *HAVCR2* (**d**) CREs in donor ND539. Each dot represents an individual sgRNA, plotted by CRISPR score and genomic position. Negative control sgRNAs as well as those targeting TSS and exons were pseudo-mapped with a 1-bp interval. **e**, Representative flow plots of PD-1 expression changes by targeting PES1, PES4, PES5 and PES8 using high scoring sgRNAs. **f**, Flow cytometry plots of the reduction in PD-1 expression in CD8 T cells from two donors by targeting PES2 and PES9. **g**, CRIPSRi mediated TIM-3 reduction by targeting the *STAT5B* TSS. Numbers on the bottom left indicate the MFI of each curve, color coding matches the corresponding curves. **h**, Quantification of TIM3 MFI by targeting *STAT5B* from (g) (mean ± s.e.m.; n = 3 donors, three independent experiments. *P<0.05, two-tailed Student’s t-test).

**Extended Data Fig. 8:**
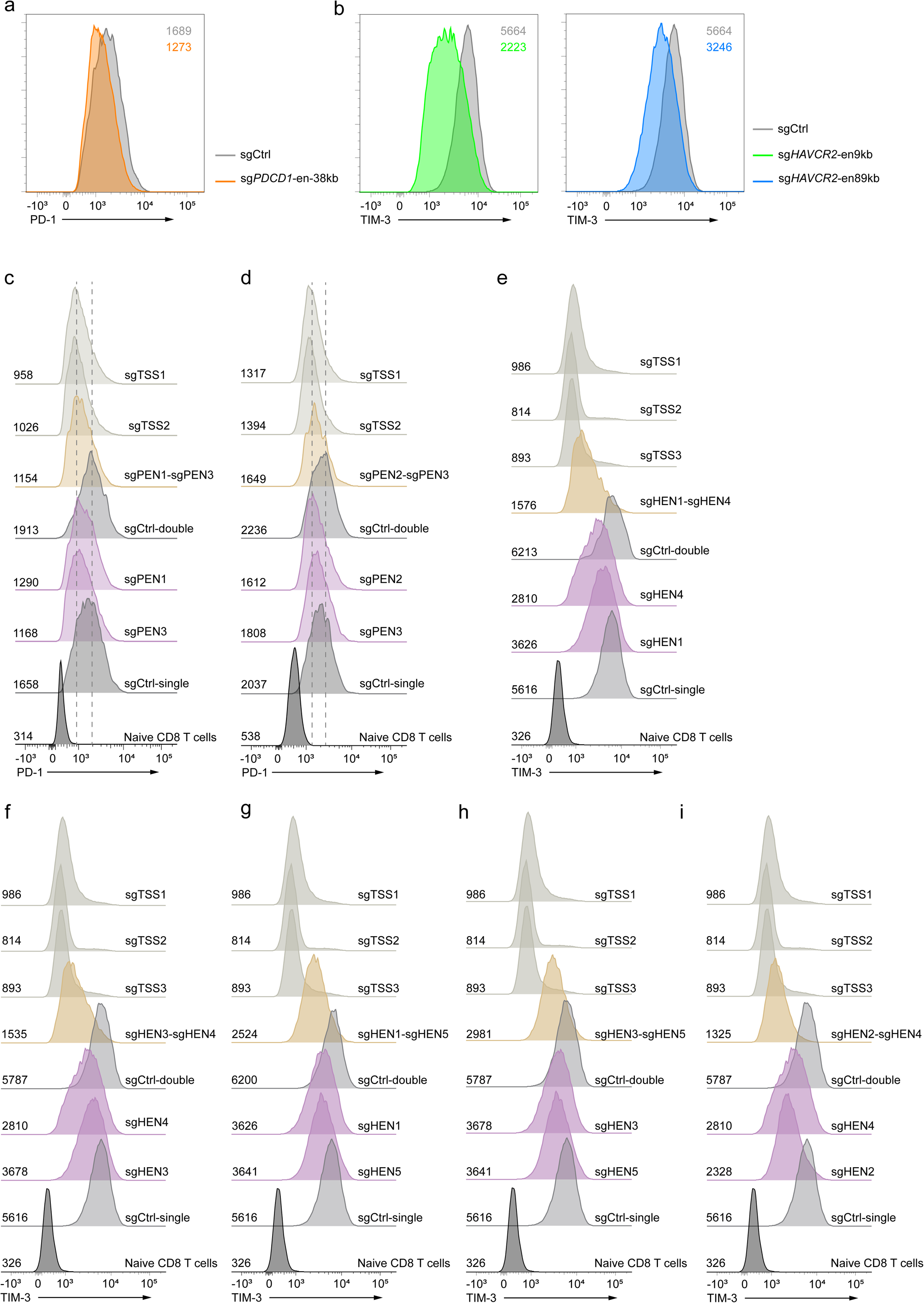
Combinatorial CRE perturbation more effectively fine-tunes IR expression. a, **b**, Flow cytometry plots of CRISPRi-mediated reduction in IR expression by targeting the *PDCD1* −38 kb CRE (**a**), or the *HAVCR2* +9 kb and +89 kb CREs (**b**). **c,d**, Representative flow cytometry plots showing PD-1 expression changes following combination CRISPRi targeting of PEN1–PEN3 (**c**) or PEN2–PEN3 (**d**) as well as single-CRE perturbation. **e–i**, Representative flow cytometry plots show TIM-3 expression after targeting specific combinations of double *HAVCR2* CRE perturbations. Numbers on the bottom left indicate the MFI of each correspondence flow cytometry plot.

**Extended Data Fig. 9:**
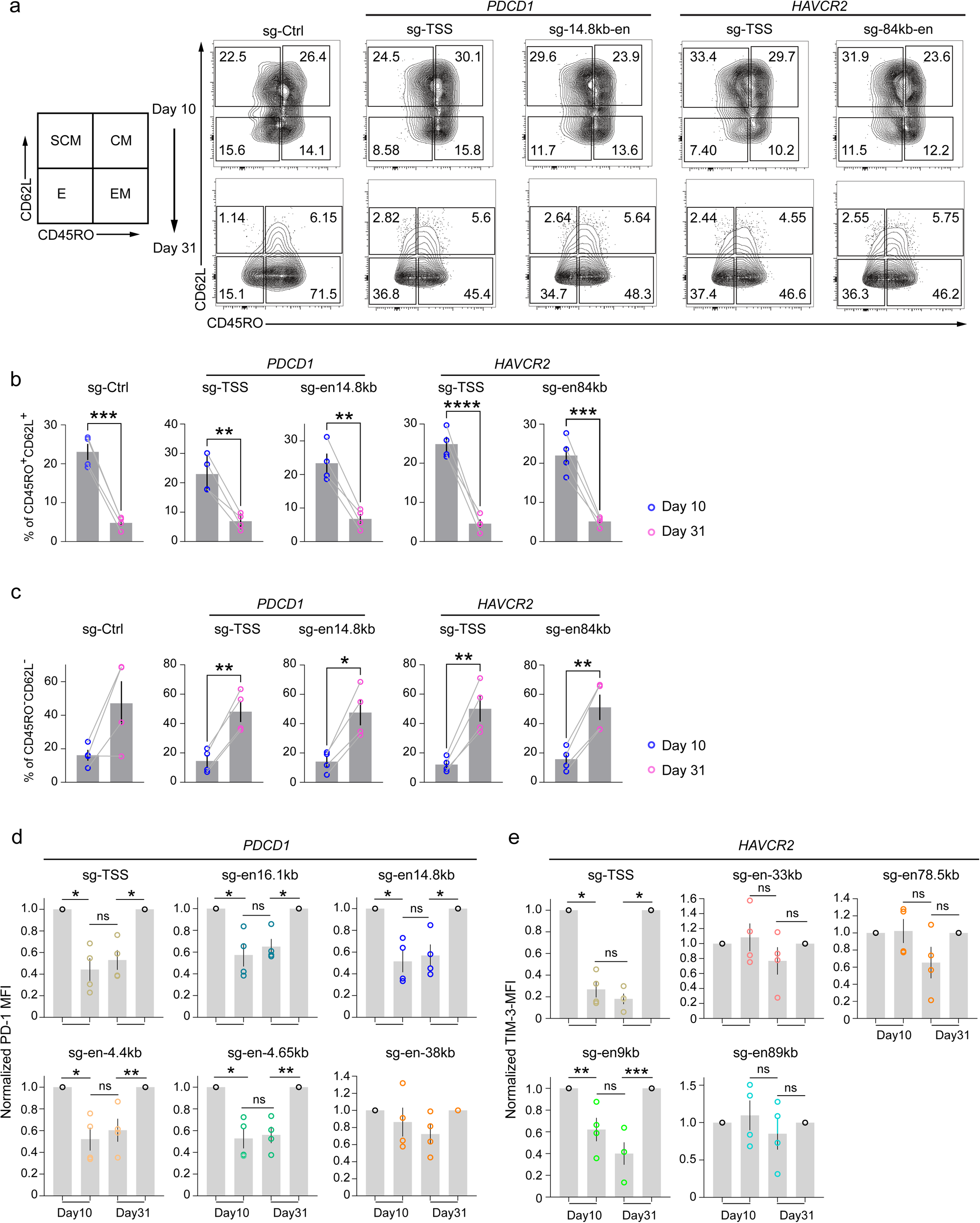
Altered differentiation state during in vitro CAR T dysfunction. **a**, Flow cytometry plots show the proportion of different subsets defined by the expression of CD45RO and CD62L in CD8 CAR T cells from CRISPRi targeted groups. **b,c**, Quantification of CD45RO^+^CD62L^+^(**b**) and CD45RO^-^CD62L^-^ subsets (**c**) in CD8 CAR T cells from different edited groups. **d,e,** Comparison of PD-1 (**d**) or TIM-3 (**e**) expression at days 10 and 31 following perturbation of TSS and CREs. b-e, Bar represents mean ± s.e.m.; n=4 donors from four different experiments. *P<0.05, **P < 0.01, ***P < 0.001 and ****P<0.0001, two-tailed Student’s t-test.

**Extended Data Fig. 10:**
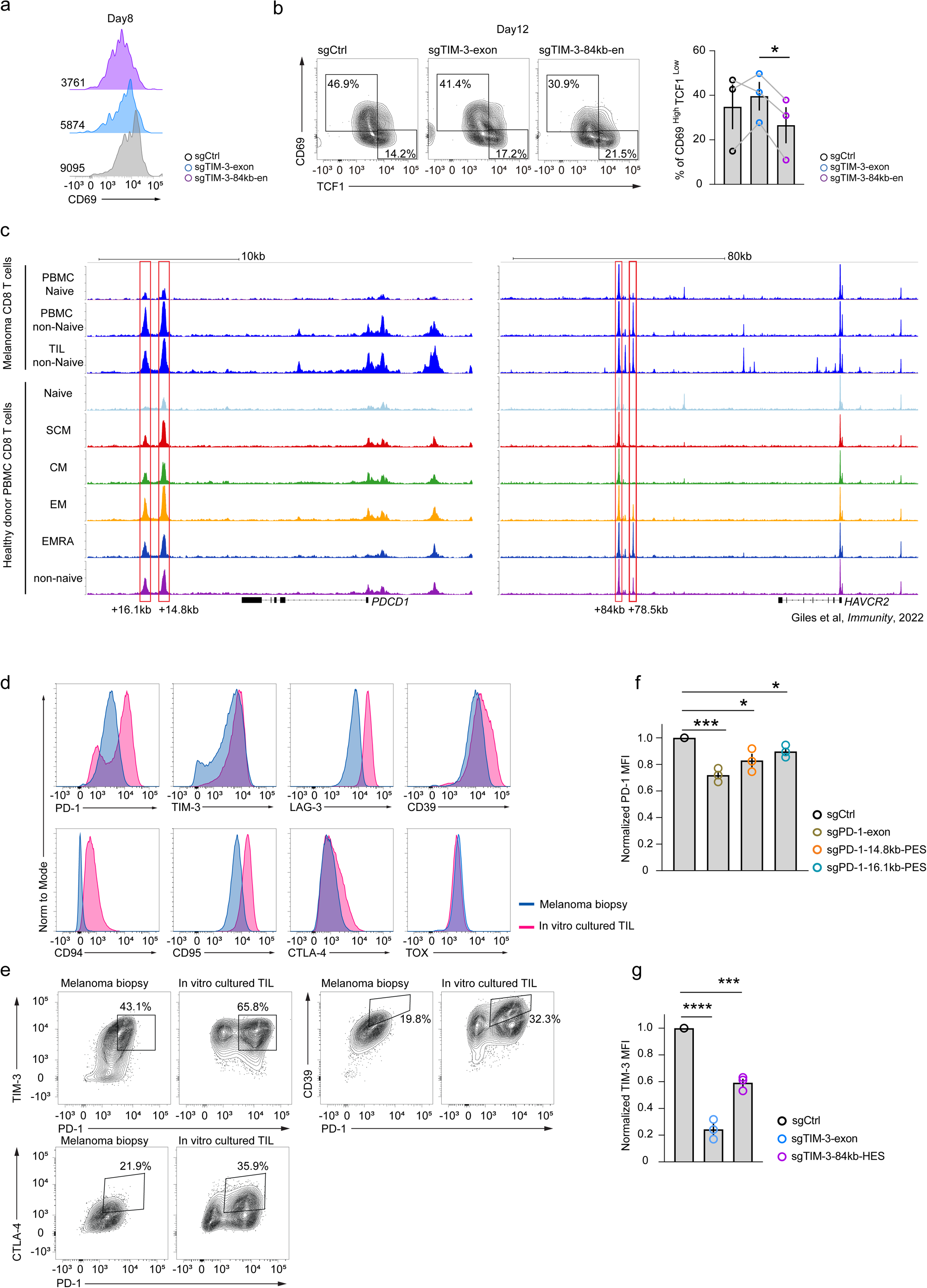
Exhaustion signature genes expression in anti-GD2 CAR T and melanoma TIL-derived CD8 T cells. a, Flow cytometry plots for CD69 expression at day 8 (Numbers on the bottom left are the MFI of each curve). **b,** Flow cytometry plots showing the proportion of the CD69^High^TCF1^Low^ subset in different anti-GD2 CAR T groups at day12. Histogram shows the quantification of results from 3 donors (mean ± s.e.m.) from a single experiment. **c,** ATAC-seq tracks showing the ACRs of *PDCD1* and *HAVCR2* loci in CD8 T cells from melanoma TIL and different subsets from healthy donor PBMCs. Red boxes indicate the *PDCD1* +14.8kb and +16.1kb CREs as well as the *HAVCR2* +78.5kb and +84kb CREs. **d,e,** Comparison of T cell exhaustion signature gene expression in CD8 T cells from melanoma biopsy and in vitro cultured TILs. **f,g**, Quantification of PD-1 (**f**) or TIM-3 (**g**) expression after targeting *PDCD1* +14.8kb and 16.1kb or *HAVCR2* +84kb CREs in in vitro cultured TIL-derived CD8 T cells. Bars represent mean ± s.e.m.; n= 3 donors, three independent experiments. *P < 0.05, ***P < 0.001 and ****P<0.0001, two-tailed Student’s t-test.

## Method

### Human PBMC and melanoma tissues

Healthy human peripheral blood mononuclear cells (PBMCs) were obtained from Human Immunology Core (HIC) at the University of Pennsylvania. Patients provided informed consent for blood and tissue collection under protocol UPCC 08607, approved by the University of Pennsylvania and the Institutional Review Board.

### Design of tile screening libraries

For each gene of interest, we identified the genomic regions of interest (ROI) as 100 kb upstream and downstream of TSS (UCSC human genome assembly, GRCh37/hg19). Repeat elements within the ROIs of *CXCR3, PDCD1* and *HAVCR2* were excluded prior to library design. Then, all possible single-guide RNAs (sgRNAs) with NGG protospacer adjacent motif (PAM) within each ROI were identified. sgRNAs containing BsmBI recognition sites or poly-T (TTTT) motifs were removed to facilitate pooled library cloning and prevent early termination of Pol III transcription of the sgRNA. The ROI was divided into 200-bp bins, the sgRNA with the least off-target potential based on previously described design principle^1^ within each bin was selected for inclusion in the tiling library. Positive control sgRNAs targeting predicted TSS of *PDCD1* or *HAVCR2* were selected from Dolcetto A and B libraries^2^. In the *CXCR3* tiling library, previously validated enhancer-targeting sgRNAs were also incorporated as positive controls^36^. For negative controls, 30 non-targeting sgRNAs from Dolcetto A and B were included, along with 60 sgRNAs targeting chromatin closed regions on chromosomes 2, 12, and X, chromosome coordinates of these regions can be found in **Supplementary Table 1**. Detailed composition of each tiling library is provided in **Supplementary Table 2-4**.

### Design of ACR screening library

Accessible chromatin regions (ACRs) were determined from previously published^32^ and newly generated ATAC-seq data in this study. The chromosome coordinates for each ACR can be found in **Supplementary Table 1**. All possible sgRNAs within each ACR were first identified, low-quality and non-functional sgRNAs were excluded using the criteria described in the tiling libraries. To construct the customized ACR-targeting library for individual genes, 4–6 sgRNAs with the least potential for off-target effects, based on previously described design principles^1^, were selected for each ACR. Positive and negative control sgRNAs were identical to those used in the tiling libraries. The ACR library for each gene can be found in **Supplementary Table 2-5**.

### Design of essential sequence screening library

All candidate sgRNAs targeting the *PDCD1* +14.8 kb and +16.1 kb enhancers, the *HAVCR2* +2.2 kb, +78.5 kb, and +84 kb enhancers were selected from their respective ACR-targeting libraries described above. For each library, sgRNAs targeting exons and regions 1kb upstream of TSS were included as positive controls. Negative controls comprised sgRNAs targeting ATAC-seq peak free regions as mentioned above, as well as non-targeting sgRNAs used in the enhancer screening libraries. Additionally, each identified enhancer was extended by 200 bp on both ends and sgRNAs targeting these flanking regions were also included as negative controls. The essential sequence screening libraries can be found in **Supplementary Table 6**.

### Construction of all-in-one vectors

To generate all-in-one vectors encoding sgRNA-U6-SFFV-KRAB-dCas9-GFP (pPW10), EFS promoter in pED9x (Addgene #163956) was first replaced with SFFV promoter, followed by replacement of mCherry marker with GFP, the insertion of PCR fragment encoding U6-sgRNA cassette with the bidirentiola U6 promoter and SFFV promoter facing the opposition direction as illustrated in **Extended Data Fig. 1a**. To generate all-in-one CRISPRi vectors with KRAB variants, site-directed mutagenesis was then performed on pPW10 to introduce double or triple amino acid substitutions at the seventh or eighth position of the ZNF10KRAB domain, yielding sgRNA-U6-SFFV-AW7EEKRAB-dCas9 (pPW11) and sgRNA-U6-SFFV-WSR8EEEKRAB-dCas9 (pPW12), respectively. For the construction of sgRNA-U6-SFFV-ZIM3KRAB-dCas9 (pPW13), the ZIM3KRAB domain from pLX303-ZIM3-KRAB-dCas9 (Addgene, #154472) was cloned into pPW10 to replace the ZNF10KRAB. GFP in pPW13 was then replaced with Thy1.1 to generate pPW18, which was used for *TBX21* enhancer screening. Thy1.1 DNA fragment was synthesized by IDT. Vector just encoding SFFV-ZIM3KRAB-dCas9 (pPW2) was constructed by replacing the EFS promoter and ZNF10KRAB in pED9x with SFFV and ZIM3KRAB. All-in-one CRISPR vector (pPW19) was constructed by replacing EFS promoter with SFFV. All vectors were constructed using In-Fusion cloning strategy (Takara Bio, 638933).

### Construction of pooled sgRNA libraries

For pooled library generation, a 24 bp forward cloning primer followed by the sequence *cgtctcacaccg* was appended to the 5′ end of each sgRNA, the 3′ end was appended with the sequence *gttttgagacg* followed by a 24 bp reverse cloning primer. Primer sequences are listed in the **Supplementary Table 8**. sgRNA libraries were synthesized as single-stranded oligonucleotide pools (Twist Bioscience). Libraries (5–10 ng) were PCR-amplified using the Q5 Hot Start High-Fidelity 2X Master Mix (NEB, M0536L), amplified products were gel purified.

For cloning, 100 ng of the amplified library was ligated with 500 ng of BsmBI-digested all-in-one CRISPRi/CRISPR vector via Golden Gate assembly in a 50 µl reaction containing 1 mM DTT (Invitrogen), 1 mM ATP (Invitrogen), 1 µl Esp3I (Thermo Fisher, ER0452), and 1 µl T7 DNA ligase (Enzymatics, L6020L). Ligation products were purified using the MinElute Enzyme Reaction Cleanup Kit (Qiagen, 28204). 2 µl of the purified product was electroporated into 30 µl of MegaX DH10B T1R Electrocomp Cells (Invitrogen, C640003). Each sgRNA was in enough replicates and produced 1000 clones in LB agar plates. Libraries were then maxi prepped (Invitrogen, K210007). The pooled sgRNAs were subjected to Sanger sequencing and deep sequencing to evaluate the accuracy and representation of the sgRNA library before genetic screening.

### Reporter assay vector

To construct reporter assay vectors, the EFS promoter upstream of GFP in the LRG2.1T vector (Addgene #108098) was replaced by a minimal TATA box promoter. Sequences of *PDCD1*, *HAVCR2* and *TBX21* enhancers were PCR amplified from genomic DNA isolated from primary human CD8 T cells. Neutral sequence (negative control) from a previously published study was synthesized by TWIST Bioscience^3^. Enhancers and negative control sequences were inserted approximately 2kb upstream of the TATA box in the reporter vector. TATA box promoter sequence used for reporter vector construction is provided in **Supplementary Table 8**.

### Lentivirus production

HEK293T cells were maintained at a confluency of 60%-70% in RPMI1640 medium supplemented with 10% Fetal Bovine Serum (Hyclone, S11150) and 100U/mL penicillin/streptomycin (Gibco, 15140122). For virus packaging, 1 × 10⁷ HEK293T cells were seeded into a 10-cm dish 15–17 h prior to transfection to ensure 90–95% confluency at the time of transfection. 80μl Polyethylenimine (Sigma, 408719) was added to 500μl OPTI-MEM medium (Gibco, 51985034), 10μg of the sgRNA-containing all-in-one vector or pooled sgRNA library was mixed with 5μg of VSVG and 7.5μg of psPAX2 in another 500μl OPTI-MEM medium, components were then combined and incubated at room temperature for 18mins. The transfection mixture was then added dropwise to the HEK293T cells. Six hours after transfection, media was replaced with 5ml fresh CTS OpTmizer T Cell Expansion media (Gibco, A1048501) supplemented with 1x non-essential amino acids (Gibco, 11140050), 10mM HEPES (Gibco, 15630080),100U/mL penicillin/streptomycin (Gibco, 15140122) and 1x viral boost (Alstem, VB100). Virus supernatant was harvested in the morning and afternoon over the following two days. After the final collection, virus was spin down at 3000rpm for 5min at 4°C and transferred to a new 50ml tube. Lentivirus was kept at 4°C for short term and stored at -80°C for long term use. HEK 293T cells were kindly provided by Joseph A. Fraietta’s laboratory (University of Pennsylvania).

### In vitro culture of human primary CD8 T cells

Primary human CD8 T cells were isolated from cryopreserved PBMCs using negative enrichment following the manufacture’s protocol (Stemcell, 17953). CD8 T cells were cultured at 1.25x10^6^ cells/mL in a 24 well plate in 1.5mL/well of TexMACS media (Miltenyi, 13009796) supplemented with 3% human AB Serum (GeminiBio, 100-512), 1x non-essential amino acids (Gibco, 11140050), 10mM HEPES (Gibco, 15630080) and 100U/mL penicillin/streptomycin (Gibco, 15140122) and stimulated with anti-CD3/CD28 beads (Gibco, 40203D) at the ratio of 1:3 (cells:beads) with 10ng/mL human IL2 (Peprotech, 200-02), 5ng/mL human IL7 (Peprotech, 200-07) and 5ng/mL human IL15 (Peprotech, 200-15). Two days after lentivirus infection, anti-CD3/CD28 beads were removed.

### Lentivirus transduction

CD8 T cells were isolated and cultured as described above. 30 hours after activation, 1.2mL medium was removed from each well and 1.2ml of lentivirus supplemented with 10ng/mL human IL2, 5ng/mL human IL7, 5ng/mL human IL15 and 8μg/mL polybrene (Sigma, H9268) was added to each well. Cells were spin infected at 2000g for 75 min at 37°C. After infection, cells were returned to 37°C for 30mins and then 500μl fresh complete TexMACS medium was added to each well.

### CRISPRi pooled CRE screen for IRs in primary human CD8 T cells

CD8 T cells were activated as described above, then transduced with the tiled or ACR CRE screening libraries. For the *PDCD1* enhancer screen, cells were harvested on day 4 post-activation, for the *HAVCR2* enhancer screen, cells were collected on day 6. Cells were washed twice with PBS then stained with aqua viability dye (Invitrogen, L34957) diluted 1:400 in PBS for 20 min at room temperature. After two washes with staining buffer (PBS containing 3% FBS), cells were incubated with 1:100 diluted anti-human PD-1(Biolegend, 329920) or anti-human TIM3 (Biolegend, 345008) antibodies at room temperature for 45 min, cells were then washed and resuspended in TexMACs media. The top 20% and bottom 20% of cells based on the expression of PD-1 or TIM3 were sorted by FACSAriaII (BD Biosciences). A target coverage of 1,000 transduced cells per sgRNA was achieved for all libraries. Isolated cells were washed twice with PBS; cell pellets were stored at -80°C for downstream genomic DNA extraction.

### CRISPRi pooled screen for *TBX21* CREs in primary human CD8 T cells

CD8 T cells were activated as described above, then transduced with the *TBX21* ACR enhancer screening library and were harvested at day 5. Cells were washed twice with PBS, then stained with aqua viability dye (Invitrogen, L34957) diluted 1:400 in PBS for 20 min at room temperature. After two washes with staining buffer (PBS containing 3% FBS), cells were stained with anti-mouse CD90.1 (Thy1.1) antibody (BioLegend, 202528) diluted at 1:200 and incubated for 45 min at room temperature. Cells were subsequently washed and fixed by 2% paraformaldehyde (Thermo Fisher, 28908) at 4°C for 10 min, then incubated with a fix and perm buffer (eBioscience, 00-5523-00) at 4°C for 30mins. After fixation, cells were stained with anti-TBET (Biolegend, 644824) at room temperature for at least 2 hours. Following intracellular staining, cells were washed by 1:10 diluted wash buffer (eBioscience, 00-5523-00) and resuspended in complete TexMACS medium. Target cell populations were isolated using the same FACS strategy described above, based on TBET expression. Isolated cells were washed twice with PBS; cell pellets were stored in -80°C for downstream genomic DNA extraction.

### Isolate genomic DNA

For *PDCD1* and *HAVCR2* CRE screening libraries transduced cells, genomic DNA was isolated from frozen cell pellets using QIAamp DNA Micro kit (Qiagen, 56304) following the manufacturer’s instructions.

For the cells isolated from *TBX21* CRE screens, genomic DNA was isolated by PureLink Genomic DNA Mini Kit (Invitrogen, K182002). Cells were resuspended in 150μl PBS and 150μl lysis/binding buffer, the reaction was incubated at 65°C for 2 hours for cell lysis and reverse crosslinking. Then, 10μl Proteinase K was added, and cells were further incubated at 55°C overnight. After genomic DNA release, the reaction was treated with 5μl RNase at 37°C for 30mins to eliminate RNA. DNA was precipitated by the addition of 150 μl ethanol (99–100%) and purified using spin columns according to the manufacturer’s protocol.

### IR CRE essential sequences (ES) screens

ES screening libraries were transduced as described above. *PDCD1* ES library infected cells were harvested at day 4, *HAVCR2* ES library infected cells were harvested at day 6. Cells were harvested, stained, and sorted as described above. Genomic DNA was isolated as described above.

### Preparation of sequencing libraries

sgRNA sequences were amplified from the isolated genomic DNA using 10μM barcoded Mis-PE primer mix and 2X Phusion Flash High-Fidelity PCR Master Mix (Thermo Fisher Scientific, F548). PCR products were purified by QIAquick PCR Purification Kit (Qiagen, 28104) per the manufacturer’s protocol. Subsequently, 150ng barcoded and adaptor ligated DNA was used to generate the final sequencing libraries using 10μM PE5/PE7 primer mix and 2X Phusion Flash High-Fidelity PCR Master Mix. The final libraries were purified by QIAquick PCR Purification Kit per the manufacturer’s protocol. Library quality was assessed using the D1000 assay (Agilent, 5067-1504) on a 2100 Bioanalyzer system (Agilent Technologies). Libraries were sequenced on a MiSeq platform (Illumina, MS-102-3001). Primer sequences can be found in **Supplementary Table 8**.

### Individual sgRNA cloning for CRE and ES verification

Sequences of sgRNAs used to verify each enhancer and essential sequence of *PDCD1*, *HAVCR2* and *TBX21* can be found in **Supplementary Table 8**. Forward and reverse oligos (Eurofin Genomics, IDT) were annealed to double strand DNA and ligated to BsmBI digested all-in-one vectors. Ligation products were transduced to Stbl3 (Invitrogen, C737303), plasmids were mini prepped (Qiagen, 27106X4). Each sgRNA was then packaged into lentivirus and transfected to human CD8 T cells as described above. CREs targeted by each sgRNA ca be found in **Supplementary Table 7**.

### Gene activation by CRISPRa

Plasmid encoding sgRNA-PCP-P65-HSF1 cassette (Addgene, 180273) were modified by replacing the puromycin resistance cassette with Thy1.1. Lentivirus encoding sgRNAs targeting the TSS and enhancers of *TBX21* and dCas9-VP64 (Addgene, 180263) were generated and co-transduced to in vitro activated CD8 T cells as described above. Four days post transduction, Thy1.1 and Cas9 double positive cells were analyzed for the expression of TBET, GZMB and Perforin using flow cytometry. sgRNAs targeting the TSS and enhancers can be found in **Supplementary Table 8**.

Antibodies used for staining different proteins and Cas9 can be found in **Supplementary Table 9**.

### Tracking enhancer dynamics using anti-GD2 CAR T cells

Primary human CD4 and CD8 T cells were isolated from cryopreserved healthy donor PBMCs. Briefly, total T cells were first enriched using the EasySep Human T Cell Isolation Kit (Stemcell, 17951) according to the manufacturer’s instructions. CD8 T cells were subsequently isolated via positive selection using the EasySep Human CD8 T cell positive selection kit (Stemcell, 17853) per the manufacturer’s protocol. In the last step of CD8 T cells isolation, the CD4 T cells remained in the supernatant which was collected. Primary CD4 and CD8 cells were then cultured separately in vitro as described above. CD4 T cells were infected with lentivirus encoding anti-GD2 chimeric antigen receptor (CAR), CD8 T cells were co-transduced with anti-GD2 CAR and all-in-one CRISPRi construct targeting distinct enhancer or TSS by lentivirus. CD4 and CD8 T cells were then stained with anti-human CD4 or CD8 antibody and anti-antibody 14G2A (Absolute antibody, Ab02227). Anti-GD2 CAR+ CD4 T cells and anti-GD2 CAR^+^CRISPRi^+^CD8 T cells were then isolated using FACS and cocultured at 1:1 ratio. 1mM Shield-1 (MedChemExpress, HY-112210) was added to the coculture media at 1:1000 dilution every two days to maintain the expression of anti-GD2 CAR^4^. On days 10, 13, 17, 21, 25, 28 and 31 of in vitro culture, expression of PD-1 and TIM-3 in CRISPRi^+^anti-GD2^+^ CD8 T cells were analyzed by flow cytometry as mentioned above. A list of antibodies can be found in **Supplementary Table 9**. The pELNS-14G2a-28HTM-28Z-FKBP plasmid used for anti-GD2 CAR generation was a kind gift from Evan Weber’s laboratory (University of Pennsylvania).

### ATAC-seq

Cryopreserved human PBMCs were thawed in EasySep buffer and then stained with aqua viability dye for 20 mins at room temperature. To sort different T cell subtypes, cells were further stained with antibodies against CCR7 (Biolegend, 353203), CD27 (Biolegend, 302832), CD45RA (Biolegend, 304134), CD49D (BD Biosciences, 566134) and CD95 (Biolegend, 305634) for 45 min at room temperature. A total of 40,000 cells were sorted and processed for ATAC-seq. Nuclei were isolated by lysing cells in a buffer containing 10mM Tris-HCl, 10mM NaCl, 3mM MgCl_2_, and 0.1% Tween-20. Immediately following isolation, nuclei were subjected to DNA tagmentation by 50μl Tn5 transposase reaction mix (Illumina, 20034197) at 37°C for 30 min. Fragmented DNA was purified by Qiagen MinElute Enzyme Reaction Cleanup Kit (Qiagen, 28204) per the manufacture’s protocol. ATAC-seq libraries were prepared by PCR amplification of DNA fragments using Nextera Index Kit (Illumina, FC-121-1012) and NEBNext High Fidelity PCR Mix (NEB, M0541L). Library fragments size distribution was measured using D1000 assay (Agilent Technologies, 5067-5582, 5067-5583) on a 2200 TapeStation (Agilent Technologies, G2965A). Libraries were quantified using the KAPA Library Quantification Kit (Roche, KK4824) and sequenced on a NextSeq 550 platform (Illumina, 20024904).

### Quantitative Real Time PCR (qRT-PCR)

Total RNA was isolated using RNeasy Plus Mini Kit (Qiagen, 74134) per the manufacturer’s protocol, cDNA was generated by RevertAid RT Reverse Transcription Kit (Thermo Fisher Scientific, K1691) following the manufacturer’s instructions. For quantitative PCR, 2μl of cDNA was combined with 1 μl RT primer mix and 5μl SYBR green master mix (Thermo Fisher Scientific, A25742) to a volume of 10μl. Reactions were run on a QuantStudio 6 Flex Real-Time PCR System (Thermo Fisher Scientific). RT primers were synthesized by IDT, sequences can be found in **Supplementary Table 8**.

### Antibody staining and flow cytometry

To analyze the expression of surface proteins, cells were washed twice with PBS then stained with aqua viability dye (Invitrogen, L34957) diluted 1:400 in PBS for 20 min at room temperature to stain live/dead cells. Then, after two washes with staining buffer (PBS containing 3% FBS), cells were resuspended in staining buffer containing surface antibody and incubated at room temperature for 45 min. cells were then washed and resuspended in TexMACs media. To analyze the expression intracellular proteins, after staining of surface proteins, cells were subsequently washed and fixed by 2% paraformaldehyde (Thermo Fisher, 28908) at 4°C for 10 min, then incubated with a fix and perm buffer (eBioscience, 00-5523-00) at 4°C for another 30 mins. After fixation, cells were resuspended in 1:10 diluted wash buffer (eBioscience, 00-5523-00) containing intracellular antibody at room temperature for at least 2 hours. Following intracellular staining, cells were washed by wash buffer and resuspended in complete TexMACS medium. All stains were performed in 50 μl volume and incubated in the dark, all washes were performed in 250-300 μl volume. A list of antibodies used in this study can be found in **Supplementary Table 9**.

Flow cytometry data were acquired on LSR II (BD Bioscience), FACSymphony™ A3 and A5 (BD Bioscience). Ultracomp eBeads (Thermo Fisher Scientific, 01-2222-42) were used for compensation. Flow cytometry data were analyzed using FlowJo (version 10.8.1, TreeStar).

### Co-culture of anti-GD2 CAR T and cancer cells

Primary human CD8 T cells were isolated and co-transduced with anti-GD2 CAR and an all-in-one CRISPR construct (sgCtrl, sg*HAVCR2*-exon or sg*HAVCR2*-84kb-en) by lentivirus to generate a negative control, TIM-3 KO or KD CAR T cells. These cells were then cocultured with 143B osteosarcoma cells at E:T ratio of 1:1. 1mM Shield-1 was added to the coculture media at 1:1000 dilution every other day to maintain the expression of anti-GD2 CAR. Cells and supernatants were collected every 3-4 days for cell number counting, CAR T cell staining and cytokine detection. 143B and T cell numbers were counted using CountBright™ Absolute Counting Beads (Thermofisher, C36950) by flow cytometry. 143B osteosarcoma cells expressing GFP and luciferase were kindly provided by Evan Weber’s laboratory (University of Pennsylvania).

### In vitro culture of human melanoma TIL

Cryopreserved single-cell suspensions from human melanoma biopsy samples were thawed and cultured in X-VIVO medium (Lonza, 02-060Q) supplemented with 30% human AB serum, 5000U human IL2, 5ng/mL IL7 and 5ng/mL IL15 for one week. Then, TILs were then co-cultured with irradiated human PBMCs from a healthy donor at the ratio of 1:5 for further expansion in the media containing 25ul/ml ImmunoCult Human CD3/CD28/CD2 T cell activator (Stemcell Technologies, 17970) and IL2, IL7 and IL15 for one week, then further expanded in X-VIVO medium containing 1000U human IL2, 5ng/mL IL7 and 5ng/mL IL15 for another two weeks.

### Electroporation of CRISPR RNPs

Lyophilized sgRNA (IDT) was resuspended in Nuclease-Free Duplex Buffer (IDT, 1072570) at 120 μM. For RNP complex formation, Cas9 nuclease (IDT, 1081058) and sgRNA were mixed at a 1:10 molar ratio and incubated at room temperature for 20 min. TIL-derived T cells were washed once with PBS, resuspended in P3 buffer (Lonza, V4XP-3032) at 1.2 × 10⁶ cells/20 μl, and combined with the RNP mixture. The cell–RNP suspension was transferred to a 16-well cuvette (Lonza, V4XP-3032) and electroporated using program EH115 on a Lonza 4D-Nucleofector® X Unit. Immediately after electroporation, 100 μl of pre-warmed X-VIVO medium was added to each well and incubated at 37 °C for 15 min before transfer to 150 μl of X-VIVO medium supplemented with IL2, IL7 and IL15 as described above.

### Luminex assay for cytokine detection

Supernatants from days 4, 8, and 12 of all coculture conditions were collected for cytokine profiling using a Luminex-based multiplex assay, performed by the Human Immunology Core at the University of Pennsylvania. Briefly, 50 µl of supernatant per well was assayed in duplicate using the ProcartaPlex Human Treg 18-Plex Panel (Thermo Fisher Scientific, EPX180-12165-901) on a FLEXMAP 3D system. Data acquisition and analysis were conducted using Luminex® xPONENT® software version 4.2 and Bio-Plex Manager™ software version 6.1.

### Screening data processing

For CRISPRi screens, raw sequencing data were first processed by demultiplexing and adaptor trimming to extract sgRNA cassette. The trimmed sequencing data were mapped to the sgRNA library without allowing mismatches, and read counts were quantified for each sgRNA. sgRNAs with <100 reads were filtered out. Read counts across all donors were normalized to the total reads of all negative controls and non-target controls. The CRISPRi score for each sgRNA were calculated by log2 fold-change (Log2FC) of normalized reads between the bottom 20% and top 20% cell populations. Reproducibility was assessed by computing the Pearson correlation of sgRNA read counts between the two donors for each screen. NCBI blast toolkit v2.10.1+ was used to align guide sequence queries. A reference index of UCSC hg19 genome was assembled using the program makeblastdb. sgRNA sequences were aligned to the reference genome using blastn with default parameters. CRISPRi scores from ACR libraries were then assigned to their correspondence sgRNAs.

For the tiled library screening data, a previously reported sliding window approach was used to calculate the average CRISPRi score of a given number of consecutive sgRNAs^7^. The number of consecutive sgRNAs (N) represents the size of sliding window. Various window sizes (N = 2, 5, 10, 15, 20, 30, 50) were selected to smooth the tiled screening data. Through assessing the correlation of CRISPRi scores between two donors under different sliding window sizes, N=20 yielded the most robust correlation (R>0.99) and was used for downstream analysis. To identify statistically significant regions, a sliding window t-test was performed, comparing the CRISPRi scores of each 20-sgRNA window to those of negative controls. Windows with Benjamini-Hochberg-corrected t-test p-values <0.05 were considered statistically significant. Statistical analyses and visualizations were performed using custom R scripts.

For data visualization, CRIPSRi scores, sliding window and accessible chromatin regions track views of ATAC-seq data were plotted using R Bioconductor package Gviz (v1.30.3). For ACR screens, CRISPRi scores were assigned to each sgRNA according to their genomic locus within the target region. For tiled screening, the smoothed CRISPRi scores from the sliding window analysis were displayed across the targeted loci.

For Cas9-indel screening, raw sequencing data was processed as mentioned above, sgRNAs with <50 reads were filtered out. Read counts across all donors were normalized to the total reads of all sgRNAs. CRISPR scores were assigned to each sgRNA according to their genomic locus within the target region. Predicted Cas9 DNA cleavage sites were mapped within each CRE to display the essential sequences distribution.

### ATAC-seq data processing

Samples were aligned to the hg19 human reference genome using Bowtie2 (2.1.0). Unmapped, unpaired, and mitochondrial reads were removed with SAMtools (1.1), ENCODE Blacklist regions were also excluded. PCR duplicates were marked and removed using Picard (1.141). Peaks were called with MACS2 (2.1.1.20160309) (FDR q < 0.001). Read counts per peak were quantified using bedtools coverage. Peaks were annotated with HOMER (4.10.3). ATAC-seq signal tracks were generated using the Gviz R package, with bigWig files normalized to library size and scaled across all samples within each dataset.

### Essential sequences and TF binding motifs analyzation

Sequences of high CRISPR score (Log_2_FC>1) sgRNAs from both donors were first mapped to the identified enhancers of *PDCD1* and *HAVCR2*. The sequences covered by all the clustered sgRNAs were termed essential sequences (ES) of an enhancer. Motifs of these ESs were identified using FIMO (MEME suites v5.1.0) with default parameters annotated using the HOMER motif database.

